# CellSIUS provides sensitive and specific detection of rare cell populations from complex single cell RNA-seq data

**DOI:** 10.1101/514950

**Authors:** Rebekka Wegmann, Marilisa Neri, Sven Schuierer, Bilada Bilican, Huyen Hartkopf, Florian Nigsch, Felipa Mapa, Annick Waldt, Rachel Cuttat, Max R. Salick, Joe Raymond, Ajamete Kaykas, Guglielmo Roma, Caroline Gubser Keller

## Abstract

Comprehensive benchmarking of computational methods for single-cell RNA sequencing (scRNA-seq) analysis is scarce. Using a modular workflow and a large dataset with known cell composition, we benchmarked feature selection and clustering methodologies for scRNA-seq data. Results highlighted a methodology gap for rare cell population identification for which we developed CellSIUS (**<u>Cell S</u>**ubtype Identification from **<u>U</u>**pregulated gene **<u>S</u>**ets). CellSIUS outperformed existing approaches, enabled the identification of rare cell populations and, in contrast to other methods, simultaneously revealed transcriptomic signatures indicative of the rare cells’ function. We exemplified the use of our workflow and CellSIUS for the characterization of a human pluripotent cell 3D spheroid differentiation protocol recapitulating deep-layer corticogenesis *in vitro*. Results revealed lineage bifurcation between Cajal-Retzius cells and layer V/VI neurons as well as rare cell populations that differ by migratory, metabolic, or cell cycle status, including a choroid plexus neuroepithelial subgroup, revealing previously unrecognized complexity in human stem cell-derived cellular populations.

## Introduction

Single-cell RNA sequencing (scRNA-seq) enables genome-wide mRNA expression profiling with single cell granularity. With recent technological advances [1,2] and the rise of fully commercialized systems [3], throughput and availability of this technology are increasing at a fast pace [4]. Evolving from the first scRNA-seq dataset measuring gene expression from a single mouse blastomere in 2009 [5], scRNA-seq datasets now typically include expression profiles of thousands [1–3] to over one hundred thousand cells [6,7]. One of the main applications of scRNA-seq uncovering and characterizing novel and/or rare cell types from complex tissue in health and disease [8–15].

From an analytical point of view, the high dimensionality and complexity of scRNA-seq data pose significant challenges. Following the platform development, a multitude of computational approaches for the analysis of scRNA-seq data emerged. These comprise tools for cell-centric analyses, such as unsupervised clustering for cell type identification [16,17], analysis of developmental trajectories [18,19] or identification of rare cell populations [9,10,20], as well as approaches for gene-centric analyses such as differential expression (DE) analysis [21–23]. Whereas a large number of computational methods tailored to scRNA-seq analysis are available, comprehensive benchmarking of, and performance comparisons between those, are scarce. This is mainly due to the lack of reference datasets with known cellular composition. Prior knowledge or use of synthetic data are commonly used to circumvent the problem of a missing ground truth. However, former knowledge might be incomplete or inaccurate and synthetic data do not capture all aspects of experimental biological data.

Here, we generated a benchmark dataset of ∼12 000 single cell transcriptomes from eight human cell lines to evaluate the performance of scRNA-seq feature reduction and clustering approaches using a modular, generally applicable workflow for the analysis of scRNA-seq data. Strikingly, results highlighted a methodology gap for the identification of rare cell types. To fill this gap, we developed a method which we called CellSIUS (**<u>Cell S</u>**ubtype **<u>I</u>**dentification from **<u>U</u>**pregulated gene **<u>S</u>**ets). For complex scRNA-seq datasets containing both abundant and rare cell populations, we propose a two-step approach consisting of an initial coarse clustering step followed by CellSIUS. Using synthetic and biological data, including rare cell populations (<0.16 %), we showed that CellSIUS outperforms existing algorithms in both specificity and selectivity for rare cell type and outlier genes (gene signature) identification. In addition, and in contrast to existing approaches, CellSIUS simultaneously reveals transcriptomic signatures indicative of rare cell type’s function(s).

Subsequently, we applied the workflow and our two-step clustering approach to biological data of unknown cell composition. We profiled the gene expression of 4 857 human pluripotent stem cell (hPSC) derived cortical neurons generated by a 3D spheroid differentiation protocol using morphogens. Analysis of this *in vitro* model of corticogenesis revealed distinct progenitor, neuronal and glial populations consistent with developing human telencephalon. Trajectory analysis identified a lineage bifurcation point between Cajal-Retzius cells and layer V/VI cortical neurons, which was not clearly demonstrated in other *in vitro* hPSC models of corticogenesis [24–27]. In addition, CellSIUS revealed rare cell populations that differ by migratory, metabolic, or cell cycle status, including a rare choroid plexus (CP) lineage, for which we experimentally validated the expression of the identified cell subtype markers at the protein level. Therefore, scRNA-seq in combination with CellSIUS provided an unprecedented resolution in the transcriptional analysis of developmental trajectories, revealed previously unrecognized complexities in human stem cell-derived cellular populations, identified rare cell populations and provided the means to isolate and characterize CP neuroepithelia *in vitro* to study neurological disorders.

## Results

### Benchmarking of feature selection and clustering approaches for scRNA-seq data reveals a methodology gap for the detection of rare cell populations

To perform a comprehensive assessment and comparison of the most recent feature selection and clustering methodologies for scRNA-seq data, we generated a scRNA-seq dataset with known cellular composition generated from mixtures of eight human cell lines. To this end, a total of ∼12 000 cells from eight human cell lines (Table 1: A549, H1437, HCT116, HEK293, IMR90, Jurkat, K562, Ramos) were sequenced using the 10X Genomics Chromium platform [3]. Cells were processed in batches containing mixtures of two or three cell lines each. One of the cell lines was present in two separate batches and confirmed that technical batch effects were minor as compared to the biological variability (Figure 1, Table 1). To infer cell type identity, we profiled each cell line individually using bulk RNA sequencing. Correlation of the single-cell to bulk expression profiles was used for cell type assignment as described in Methods (Figure 1A- B). Cells that did not pass quality control (QC) or could not be unambiguously assigned to a cell line (614 cells, ∼5%) were discarded, leaving 11 678 cells of known cell type (Figure 1C and S1, Table 1).

**Table 1:** Cell lines and culture conditions used in this study

| Cell line | Gender | Cell type | Tissue of origin | Obtained from | Culture conditions |
| --- | --- | --- | --- | --- | --- |
| A549 | M | Alveolar basal epithelial (adherent) | Lung adenocarcinoma | ATCC CCL-185 | ATCC F12K (ATCC, P/N 30-2004) +10% FCS (AMIMED, P/N 2-01F36-I). |
| H1437 | M | Epithelial / glandular (adherent) | Lung adenocarcinoma, derived from metastatic site: pleural effusion | ATCC CRL-5872 | RPMI (Invitrogen, P/N A1049101) +10% FBS (ATCC, P/N SCRR-30-2020) |
| HCT116 | M | Epithelium-like (adherent) | Colon carcinoma | ATCC CCL-247 | ATCC McCoy's 5A (ATCC, P/N 30-2007) + 10% FCS (AMIMED, P/N 2-01F36-I) |
| HEK293 | F | adherent | Transformed cell line, derived from embryonic kidney | ATCC, P/N CRL-1573 | ATCC EMEM (ATCC, P/N 30-2003) +10% FCS (AMIMED, P/N 2-01F36-I) |
| IMR90 | F | Fibroblast (adherent) | Fetal lung | ATCC CRL-186 | ATCC EMEM (ATCC, P/N 30-2003) 10% FCS (AMIMED, P/N 2-01F36-I) |
| Jurkat | M | T-cell (suspension) | Childhood T acute lymphoblastic leukemia | ATCC, P/N TIB-152 | RPMI (Invitrogen, P/N 61870-044) + 10% FCS (AMIMED, P/N 2-01F36-I) |
| K562 | F | Undifferentiated, lymphoblast with granulocyte/erythrocyte/monocyte characteristics (suspension) | Chronic myelogenous leukemia, BCR-ABL1 positive | ATCC, P/N CRL-1573 | RPMI (Invitrogen, P/N 61870-044) + 10% FCS (AMIMED, P/N 2-01F36-I). |
| Ramos | M | B-cell (suspension) | Burkitt's lymphoma | ATCC, P/N CRL-1596 | Batch 3: RPMI (Invitrogen, P/N A1049101) +10% FBS (ATCC, P/N SCRR-30-2020)<br>Batch 4: RPMI (Invitrogen, P/N 61870-044) + 10% FCS (AMIMED, P/N 2-01F36-I) |

**Figure 1:**
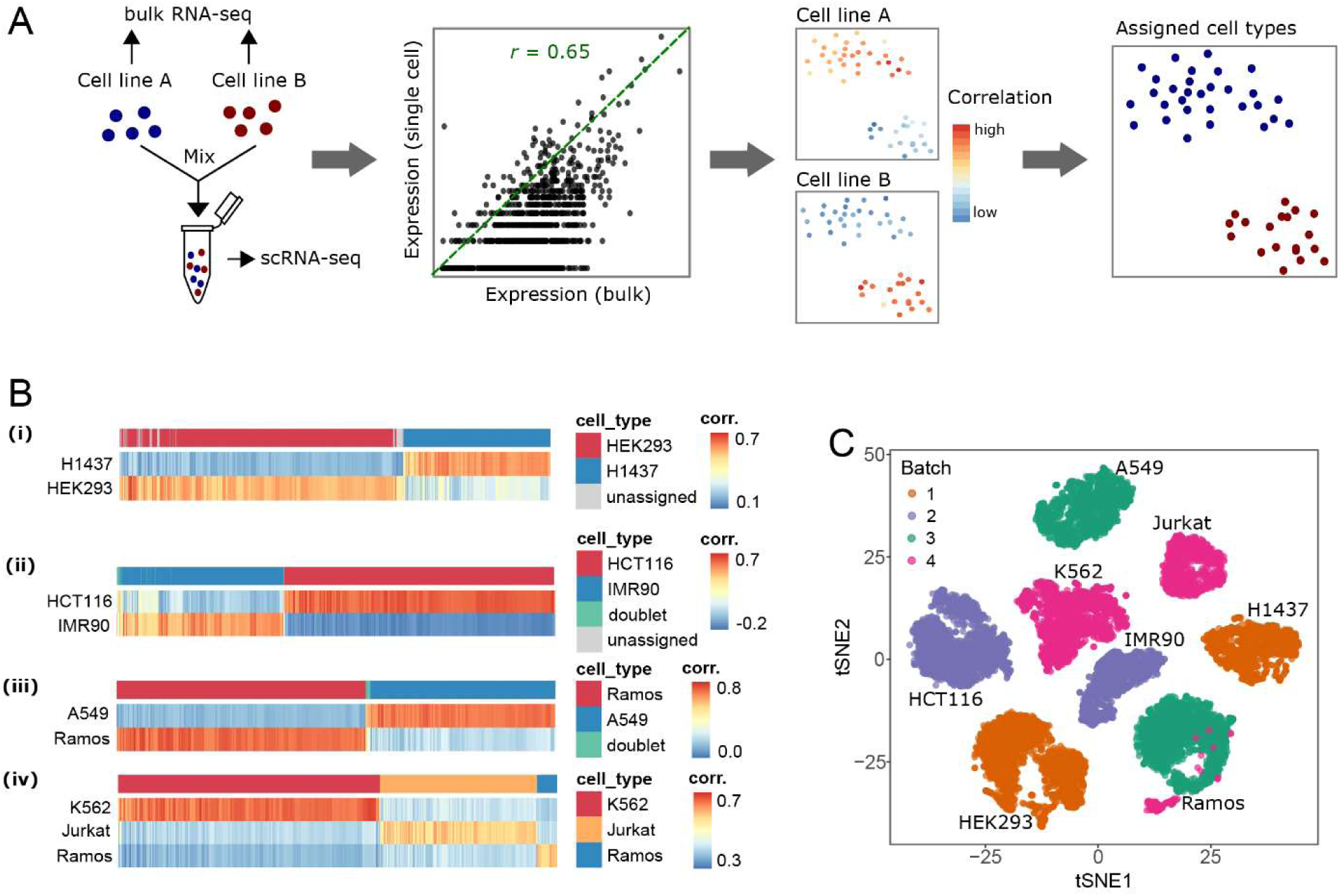
Generation of a scRNA-seq dataset with known cellular composition. A: Schematic illustration of the experimental setup. Eight human cell lines were individually profiled by bulk RNA-seq and mixed in four batches containing mixtures of two or three cell lines each for scRNA-seq profiling. Correlation of the single-cell to bulk expression profiles was used for cell type assignment as described in Methods. B: Visualization of correlations between single cell and bulk expression profiles for each batch. The top row represents cell type assignment. Single cells were assigned to the cell type correlating most with their expression profile as described in Methods. Cells with z-scored correlations below 0.2 were not assigned to any cluster. Cells that correlate strongly with more than one bulk expression profile likely represent doublets and were excluded from future analyses. C: tSNE map, colored by batch.

Using available open source tools from R [28] and Bioconductor [29], we assembled a modular workflow for the analysis of scRNA-seq data (Figure 2A). The workflow contains five modules; (i) quality control, (ii) data normalization, (iii) feature selection, (iv) clustering and, (v) identification of marker genes. Based on recent publications, the quality control and normalization modules were based on the popular scater [30] and scran [31] packages. Scran was set as the default normalization based on a recent benchmarking study by Vallejos *et al*. [32] showing that scran was superior for recovering true size factors compared to other methods. For the marker gene identification module we used the Wilcoxon test [33] by default, and provided wrappers to MAST [22] and Limma-trend [34], based on Soneson *et al*. *’s* [35] comprehensive assessment of a large number of DE analysis methods for their performance for controlling type I and type II error rates whilst being scalable to large datasets.

**Figure 2:**
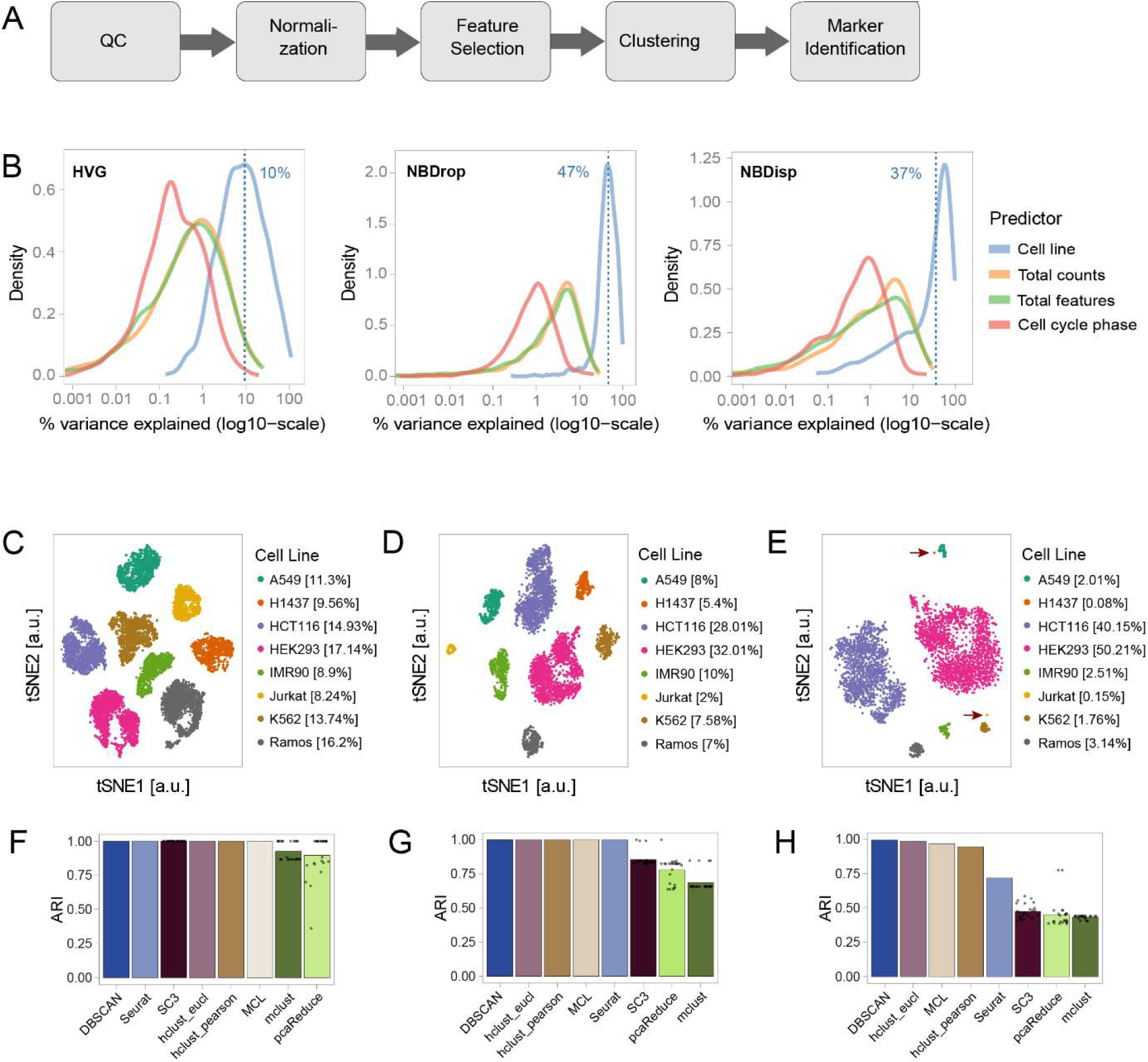
Performance assessment of feature selection and clustering methods. A: Overview of the computational analysis workflow. B: Benchmarking of feature selection methods. In each case, the top 10% of features were selected using either a mean-variance trend to find highly variable genes (HVG, left) or a depth-adjusted negative binomial model (DANB) followed by selecting genes with unexpected dropout rates (NBDrop, middle) or dispersions (NBDisp, right). Plots show the percentage of variance explained by each of the four predictors to t the total observed variance: cell line, total counts per cell, total detected features per cell and predicted cell cycle phase. The blue dashed line indicates the average for the predictor cell line. C-E: tSNE projections of the full dataset (C) and two sub-sampled datasets with unequal proportions between different cell lines (D,E). F-H: Comparison of clustering assignments by different methods on the full dataset (F), subset 1 (G) and subset 2 (H). Stochastic methods (SC3, mclust, pcaReduce) were run 25 times. Bars represent mean adjusted rand index (ARI) and dots correspond to results from individual runs. All other methods are deterministic and were run only once.

For the feature selection and clustering modules, no comprehensive method performance comparisons were available. We leveraged our dataset of known cell composition to benchmark available approaches. Briefly, we benchmarked feature selection methods using either a mean-variance trend to find highly variable genes (HVG, [36]) or a depth-adjusted negative binomial model (DANB, [37]) for selection of genes with unexpected dropout rates (NBDrop) or dispersions (NBDisp). The top 10% genes selected by HVG, NBDisp and NBDrop were included in Tables S1. Using linear modelling as implemented in the plotExplanatoryVariables function from scater [30], we quantified the influence of these feature selection methods on the contribution of four predictors to the total observed variance: cell line, total counts per cell, total detected features per cell and predicted cell cycle phase (Figure 2B). Results highlighted that: i) for HVG selected genes, cell line accounted for 10% of the total variance only; ii) for NBDisp and NBDrop selected genes, the percentage of total variance explained by cell line increased to 37% and 47%, respectively, with half of the selected features common to both methods; iii) genes selected only by NBDisp were generally low-expressed (Table S1), highlighting a drawback of variance-based feature selection [37] and; iv) NBDrop selected features showed an increased contribution of library size (i.e. total detected features and total counts per cell) to the total variance. In our dataset, the number of total features co-varied with cell type and cell cycle indicating that library size is partially dependent on the cell line (Figure S1), and thus determined by both technical and biological factors.

For the clustering module, we performed benchmarking after feature selection using NBDrop. We investigated methods (Table 2) that were developed specifically for scRNA-seq data (SC3 [16], Seurat [1], pcaReduce [17]) as well as more classical approaches (hclust[38], mclust[39], DBSCAN[40], MCL[41,42]) by *in silico* subsampling of our dataset of known composition in two subsets with different cell type proportions (later referred to as subset 1 and subset 2, Figure 2C-E, Table S2). Subset 1 consisted of 4 999 cells from eight cell types with abundance varying between 2% and 32%. Subset 2 consisted of 3 989 cells with two major cell populations including 90% of all cells of this subset, four medium to low abundant (between 1% and 5%) and two rare cell types with abundances below 1%, containing 3 (0.08%) and 6 (0.15%) cells, respectively. We applied each clustering method to the complete dataset as well as to both subsets, using principal component analysis (PCA) [43,44] to project the original expression values to vectors in a lower dimensional space and calculating all distances based on these projections. We then assessed the quality of the classification by calculating the adjusted Rand index (ARI) [45] between assignment and true cell line annotation.

**Table 2:** Overview of clustering algorithms benchmarked in this study

| Method | Unsupervised # clusters? | Input | Underlying model | Expected cluster shape and size<sup>2</sup> | Run time<sup>3</sup> |
| --- | --- | --- | --- | --- | --- |
| SC3 [16] | Yes | Normalized data as an SCESet, distances and transformations are calculated internally | K-means clustering on various distances & transformations, hierarchical clustering of consensus matrix | Spherical, equal sizes | 35 min (using hybrid SVM approach) |
| Hclust + dynamic tree cut [38,71] | No | Pearson or euclidean distance in PCA space | Agglomerative clustering | None | 1 min |
| pcaReduce [17] | No | Normalized data, PCA is performed internally | K-means + hierarchical clustering | Spherical, equal sizes | 3 min |
| Seurat [1] | Yes | Normalized, log2 transformed counts as a Seurat object | Graph based | None | 9 min |
| MCL [41,42] | Yes | Pearson distance in PCA space | Graph based | None | Build graph: >1h<br>Run MCL: 7 min |
| mclust [39] | Yes (via cross-validation or BIC) | Principal component scores | Gaussian mixture model | Ellipsoid, size can vary | 6 min |
| DBScan [40,72] | Yes | Euclidean distance in PCA space | Clusters are defined as regions of high density separated by regions of low density | None, but clusters have to be compact and clearly disconnected | 2 min |

<sup>2</sup> By size, we are referring to the actual distribution of the points in space, NOT the number of points in the cluster. For a Gaussian ellipsoid, size is parameterized by the covariance matrix.
<sup>3</sup> Run time was estimated using the system.time() function in R. The time shown here refers to the full dataset (12000 cells). Analysis was run on 64-bit Intel(R) Xeon(R) CPU E7-4850 v2 @ 2.30GHz with 1TB of RAM in R 3.4.1 under Red Hat Enterprise Linux Server release 6.9 (Santiago). SC3 was run on 8 cores, all other methods on a single core.

On the full dataset, most methods resulted in a perfect assignemnt (Figure 2F) with only two of the stochastic methods – pcaReduce and mclust – yielding an average ARI of 0.90 and 0.92. In contrast, on subset 1, where cell type proportions were no longer equal, k-means based methods and mclust failed to identify the different cell types correctly and resulted in average ARI of 0.85 (SC3), 0.78 (pcaReduce) and 0.69 (mclust) (Figure 1G). On subset 2, all methods failed to correctly identify rare (6 cells, 0.16% of total cells) cell types (Figure 1H). DBSCAN achieved the highest ARI (0.99) classifying rare cells as outliers (“border points”). All other methods merged rare cells with clusters of abundant cell types resulting in lower ARI of 0.98 (hclust on Euclidean distance), 0.96 (MCL), 0.96 (hclust on correlation distance) and 0.76 (Seurat). In conclusion, our results showed that most clustering methods performed well in identifying populations defined by more than 2% of total cells. Yet, none of the methods could identify rare populations, highlighting the need for dedicated tools tailored to detecting rare cell types.

### Development of CellSIUS for rare cell population identification and characterisation

To overcome the above-mentioned limitations, we developed a novel method to identify rare cell populations which we called CellSIUS (**<u>Cell S</u>**ubtype **<u>I</u>**dentification from **<u>U</u>**pregulated gene **<u>S</u>**ets). CellSIUS takes as input the expression values of N cells grouped into M clusters (Figure 3A). For each cluster *C*_*m*_, candidate marker genes *g*_*m1*_, *g*_*m2*_, …, *g*_*mj*_ that exhibit a bimodal distribution of expression values with a fold change above a certain threshold (fc_within) across all cells within *C*_*m*_ are identified by 1-dimensional k-means clustering (with k=2). For each candidate gene *g*_*mi*_, the mean expression in the second mode is then compared to this gene’s mean expression level outside *C*_*m*_ (fc_between), considering only cells that have non-zero expression of *g*_*mi*_ to avoid biases arising from stochastic zeroes. Only genes with significantly higher expression within the second mode of *C*_*m*_ (by default, at least a 2-fold difference in mean expression) are retained. For these remaining cluster specific candidate marker genes, gene sets with correlated expression patterns are identified using graph-based clustering. In a last step, cells within each cluster *C*_*m*_ are assigned to subgroups by 1-dimensional k-means clustering of their average expression of each gene set.

**Figure 3:**
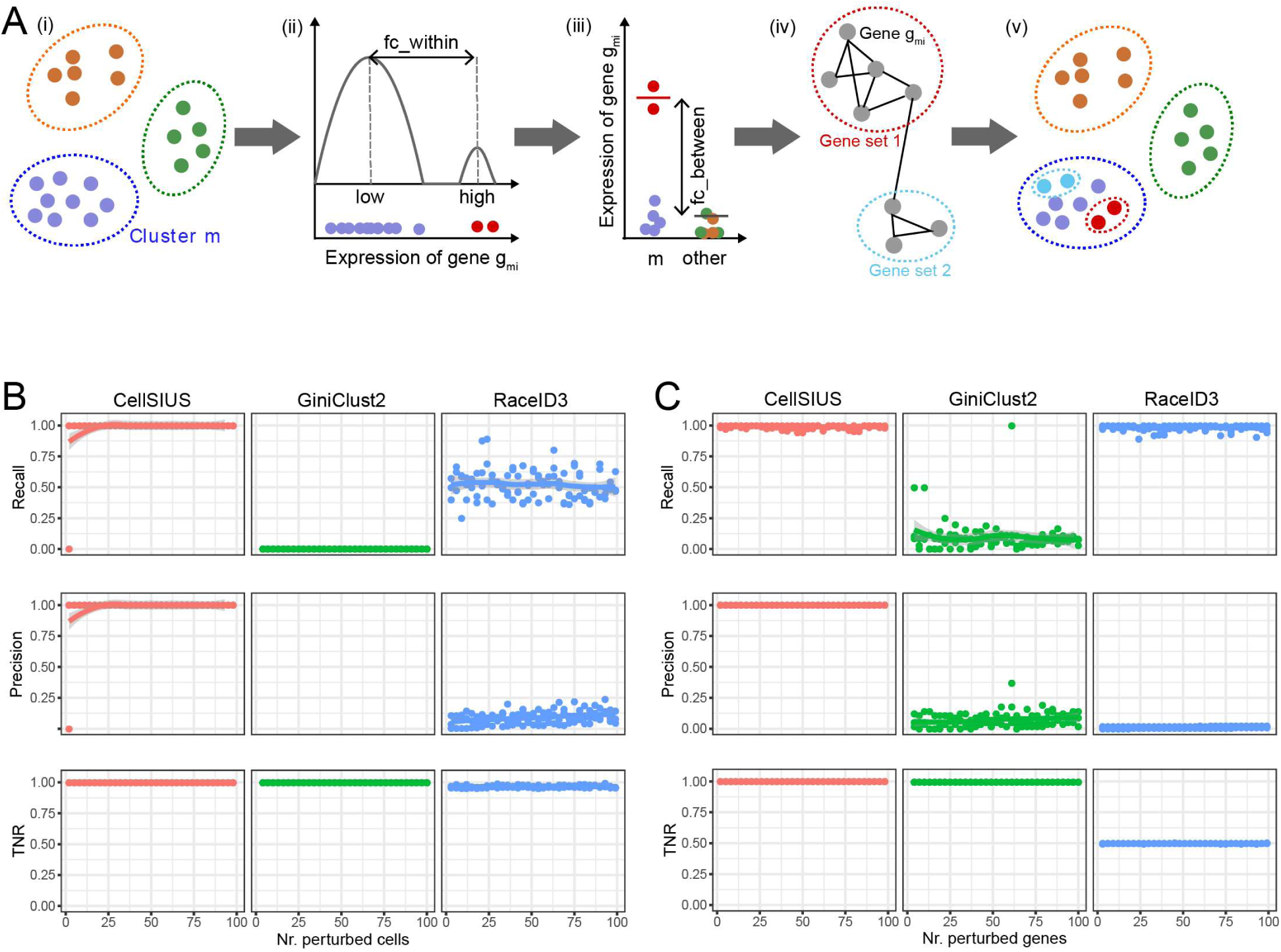
Development and benchmarking of CellSIUS. A: Schematic overview of CellSIUS. Starting from an initial assignment of N cells in M clusters (i), within each cluster, genes with a bimodal distribution are identified (ii) and only genes with cluster-specific expression are retained (iii). Among the candidate genes, sets with correlated expression patterns are identified by graph-based clustering (iv). Cells are assigned to subgroups based on their average expression of each gene set (v). B, C: Performance comparison of CellSIUS to GiniClust2 and RaceID3 in detecting cells from sub-clusters and their signatures. B: Recall, precision and true negative rate (TNR) with regards to the detection of rare cells in synthetic data when varying the number of rare cells from 2 (0.2%) to 100 (10%) C: Recall, precision and true negative rate (TNR) with regards to the detection of outlier genes (gene signature) in synthetic data when varying and the number of signature genes from 2 to 100.

The overall idea behind CellSIUS is similar to RaceID3 [46] and GiniClust2 [20], two recent methods for the identification of rare cell types in scRNA-seq datasets. All of these algorithms combine a global clustering with a second assignment method tailored to the identification of rare cell types. However, in contrast to existing methods, CellSIUS requires candidate marker genes to be cluster specific, and therefore we hypothesized that our method will be more specific and less sensitive to genes that co-vary with confounders such as the total number of detected genes per cell. To overcome biases associated to the high dropout rates in scRNA-seq, CellSIUS considers only cells that have non-zero expression for the selected marker genes. Finally, in contrast to both RaceID3 and GiniClust2, CellSIUS directly returns a gene signature for each of the new cell subpopulations recovered.

### CellSIUS outperforms existing algorithms in the identification of rare cell populations

We first compared CellSIUS performance to RaceID3 [46] and GiniClust2 [20], using a synthetic dataset. Briefly, we used the expression values of 1 000 K562 cells from our dataset to estimate the parameters for the simulation and generated two homogeneous populations of 500 cells (later referred to as Clusters 1 and 2). We confirmed the mean-variance and mean-dropout relationships, library sizes and percentage of zero counts per cells and per gene were similar to the underlying real data (Figure S2A-F). For this data, both CellSIUS and GiniClust correctly identified the two predefined clusters whereas RaceID3 detected a large number of false positives clusters by identifying outlier cells and re-assigned those to new cluster centers (Figure S2G).

To assess the specificity and sensitivity of CellSIUS for the identification of rare cell types, we simulated a series of cell compositions comprising two abundant and one increasingly rare cell types consisting of 2 to 100 cells (0.2–10% of the cluster size) that was generated by permutation of the expression values of 20 genes. To assess the response of CellSIUS’ output to parameter changes, we varied fc_within (minimum difference in log2 scale between the first and second mode of the bimodal gene expression distribution) and fc_between (minimum difference in log2 scale in gene expression between cluster-specific and other cells in the dataset). Results showed that CellSIUS only failed to detect rare cell populations consisting of 2 cells for fc_within of 1 and 2 and fc_between of 0.5 and 1 and never falsely identified rare populations (Figure S3A). We next compared the performance of CellSIUS to RaceID3 and GiniClust2 by computing (i) recall as the fraction of rare cells correctly assigned to new clusters; (ii) precision as the fraction of true rare cells among all cells not assigned to the two main clusters and (iii) true negative rate (TNR) as the fraction of abundant cells that were correctly assigned to the two main clusters. To enable a more direct comparison between the methods, benchmarking analyses were carried out with a predefined initial clustering for all approaches. CellSIUS had a recall of 1 for rare cell populations consisting of more than 2 cells. In contrast GiniClust2 did not identify any rare cell populations and RaceID3 recalled only ∼50% of true positives (Figure 3B, top panel). Additionally, CellSIUS exhibited a TNR of 1.0 and thus a precision of 1.0 (except in the one case where no true positives were recovered). Whilst GiniClust2’s TNR was also 1.0, the precision could not be defined due to the lack of identification of true and false positives. RaceID3 had a low TNR (mean=0.95, sd=0.01), resulting in low precision (mean= 0.1, sd=0.1) (Figure 3B, middle and bottom panel).

To assess the specificity and sensitivity of CellSIUS for the identification of outlier genes, we generated a second set of populations. Briefly, 20 cells (∼2% of cluster1 cells) were added by perturbing between 2 and 100 genes. We varied CellSIUS fc_within and fc_between parameters as described above. CellSIUS only failed to identify true positive genes if their expression was lower than fc_within and fc_between (see Figure S3B, top panel), and never falsely identified outlier genes (see Figure S3B, bottom panel). We next compared the performance of CellSIUS to RaceID3 and GiniClust2 by computing (i) recall; (ii) precision and (iii) TNR as above with respect to genes. In comparison to CellSIUS, GiniClust2 showed a poor performance (Figure 3C top panel), consistent with failing to detect rare cell population. In contrast, RaceID3 performed slightly better than CellSIUS in terms of recall, however, with a precision cost. Whereas both precision and TNR were 1.0 for CellSIUS, RaceID3 had a low TNR (0.5) and consequently a low precision (mean=0.012, sd=0.007) (Figure 3C, top and bottom panels). In summary, using synthetic data, we showed an increased sensitivity and specificity of our algorithm for rare cell type identification and outlier gene identification compared to GiniClust2 and RaceID3 (Figure 3B and C).

We next benchmarked CellSIUS’ specificity and selectivity using our dataset of known cell composition, randomly subsampling 100 HEK293 cells, 125 Ramos cells, and including 2, 5 or 10 Jurkat cells. Only cells assigned to be in cell cycle phase G1 were considered to ensure within-cluster homogeneity. To simulate varying degrees of transcriptional difference between the rare cell type (Jurkat) and its closest more abundant cell type (Ramos), we adapted an approach recently presented by Crow *et al*. [47]. Briefly, from the initial dataset, 25 Ramos cells were held out. Subsequently, an increasing fraction of gene expression values in the Jurkat cells were replaced by the respective values in the held out Ramos cells, thus diluting the Jurkat-specific gene expression profile and making the Jurkat cells more and more similar to Ramos (Figure 4A). Using this approach, we generated datasets with two equally sized abundant populations (HEK293 and Ramos, 100 cells each) and one rare population (Jurkat, varying between 2, 5 and 10 cells). We predefined two initial clusters: Cluster 1 contained all HEK293 cells, and cluster 2 combined Ramos and Jurkat cells.

**Figure 4:**
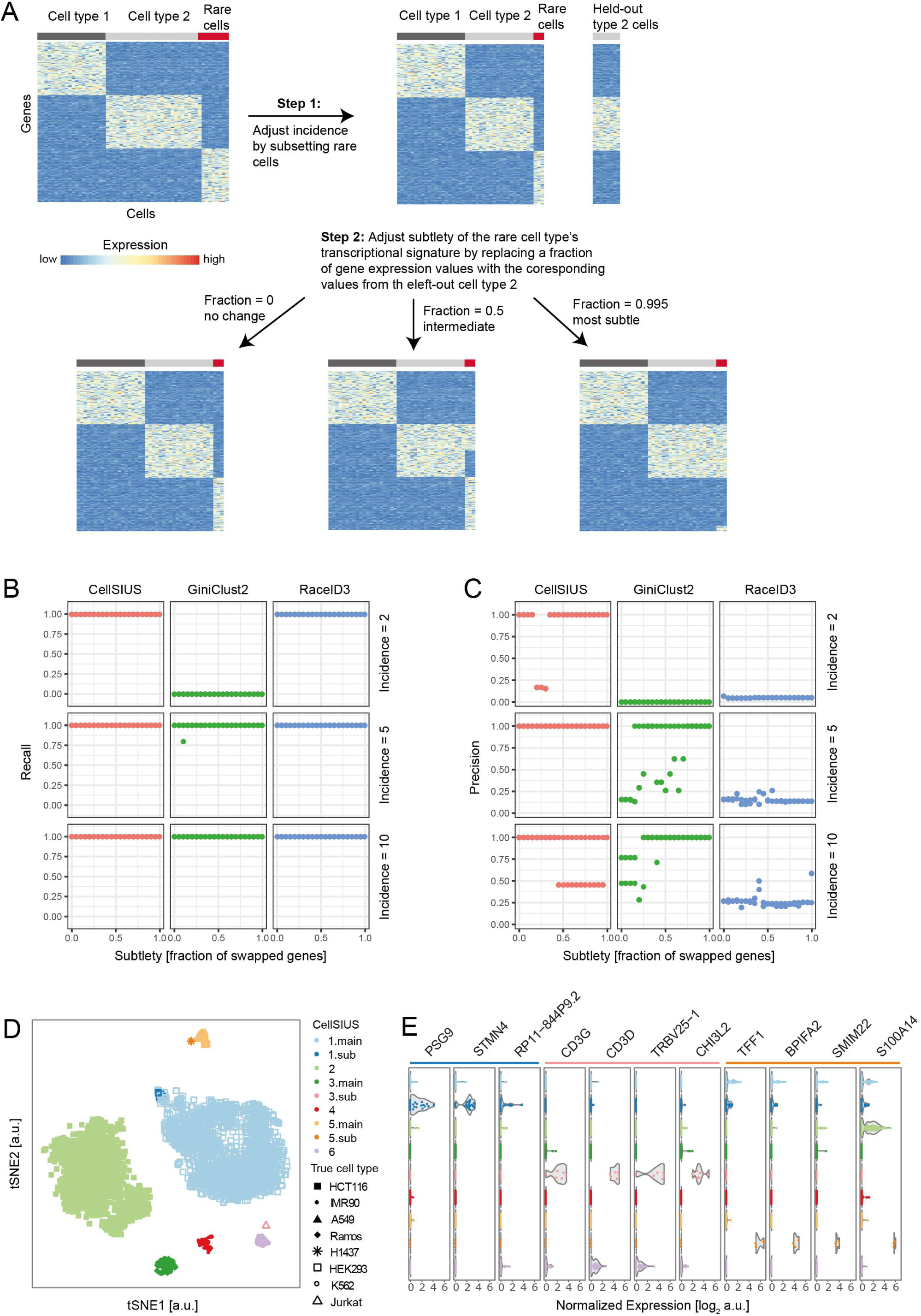
CellSIUS benchmarking on cell line data. A: Schematic overview of dataset perturbations. Starting from a dataset containing three cell types (100 HEK293 cells, 125 Ramos, 10 Jurkat), we first generated a defined number of rare cells by subsampling. In addition, we partitioned the Ramos cells in two, leaving out 25 cells from the dataset for later use. Next, we adjusted the subtlety of the transcriptional difference between the rare (Jurkat) cells and their closest neighbor (Ramos) by swapping a fraction of gene expression values in the Jurkat cells with the corresponding value in the left-out Ramos cells. We then pre-defined an initial cluster assignment as Cluster 1 = HEK293, Cluster 2 = Ramos and Jurkat and assess whether different algorithms for detecting rare cell types are able to correctly classify the Jurkat cells as rare. B, C: Comparison of CellSIUS to GiniClust2 and RaceID3 for varying incidence of the rare cell type and varying subtlety of the transcriptional signature. For each algorithm, we assessed the recall (A), i.e. the probability of detecting a rare cell type, and precision (B), i.e. the probability that a cell which is classified as rare is actually a rare cell. D: tSNE projection of subset 2 of the cell line dataset, colored by CellSIUS assignment. Cluster numbers correspond to the main clusters identified by MCL, clusters labeled x.sub indicate the CellSIUS subgroups. Symbols correspond to the cell line annotation. E: Violin plot showing the main markers identified by CellSIUS, grouped by cluster.

We then tested the ability of CellSIUS, RaceID3 and GiniClust2 to identify rare cell types for varying incidence (i.e. total number of rare cells) and subtlety (i.e. fraction of Jurkat genes replaced by Ramos genes). We assessed the recall (Figure 4B) and precision (Figure 4C) as above. Results showed a high sensitivity of all three methods for very subtle transcriptional signatures (99.5% of genes replaced, corresponding to 230 unperturbed genes) and low incidence (down to two cells except for GiniClust2). However, CellSIUS exhibited high precision (88.4% on average), in comparison to GiniClust2 (51.6% on average) and RaceID3 (15.6% on average). Having shown that CellSIUS is more sensitive and specific for the identification of rare cell types and outlier genes using synthetic and simulated biological data, we tested its ability to reveal transcriptomic signatures indicative of rare cell type’s function(s). We applied CellSIUS to subset 2 of our dataset of known composition (Table S2) with 6 clusters predefined using MCL (Figure 4D). CellSIUS identified three subgroups (Jurkat, H1437 and a small subgroup of IMR90 cells) within the 6 initial clusters characterized by upregulation of three or more genes (Figure 4E). Notably, the two strongest signatures were obtained for the two subgroups corresponding to Jurkat and H1437 cells with top marker genes consistent with previous knowledge: *CD3G* and *CD3D*, both of which are known T-cell markers [48] being the top markers for Jurkat (T-cell lymphoma) and *TFF1* and *BPIFA2*, both shown to function in the respiratory tract, [49] [50] being the top markers for H1437 (lung adenocarcinoma, epithelial/glandular cell type).

Taken together, these results show that CellSIUS outperforms existing methods in identifying rare cell populations and outlier genes from both synthetic and biological data. In addition, CellSIUS simultaneously reveals transcriptomic signatures indicative of rare cell type’s function.

### Application to hPSC-derived cortical neurons generated by 3D spheroid directed-differentiation approach

As a proof of concept, we applied our two-step approach consisting of an initial coarse clustering step followed by CellSIUS to a high quality scRNA-seq dataset of 4 857 hPSC-derived cortical neurons generated by a 3D cortical spheroid differentiation protocol with patterning factors (Figure 5A, Table S3, Methods). During this *in vitro* differentiation process, hPSCs are expected to commit to definitive neuroepithelia, restrict to dorsal telencephalic identity and generate neocortical progenitors (NP), Cajal-Retzius (CR) cells, EOMES+ intermediate progenitors (IP), layer V/VI cortical excitatory neurons (N), and outer radial-glia (oRG) (Figure 5B). Generation of layer V/VI neuronal populations was confirmed by immuno-fluorescence analysis of D86 cultures upon dissociation and plating, showing robust expression of deep-layer cortical neuronal markers TBR1 and CTIP2 (Figure 5C).

**Figure 5:**
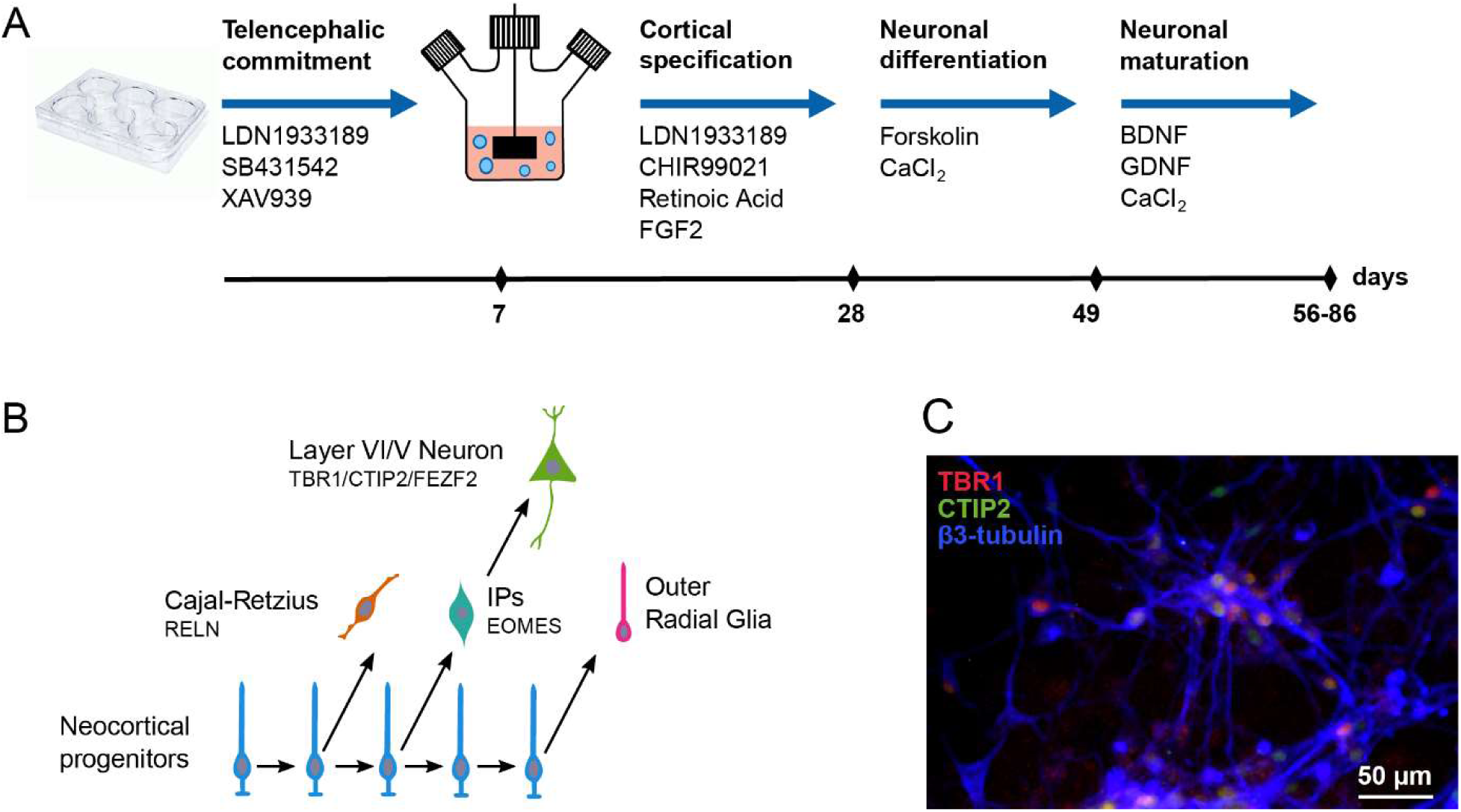
In vitro differentiation of hPSCs into cortical excitatort neurons. A: Schematic overview of the 3D cortical spheroid differentiation protocol. hPSCs grown as a monolayer were patterned to telencephalon and differentiated in suspension culture by stage-specific application of small molecules. B: Illustration of neurogenesis. After committing to definitive neuroepithelia and restricting to dorsal telencephalic identity, hPSCs generate neocortical progenitors which further give rise to Cajal-Retzius (CR) cells, EOMES+ intermediate progenitors (IPs), layer VI and V cortical excitatory neurons (N) and outer radial glia (oRG). C: Immunofluorescence confirms the robust expression of deep-layer cortical neuronal markers (TBR1, CTIP2) in hPSC derived neurons (β 3-tubulin).

Cortical neurons generated by the 3D spheroid protocol co-cultured with rat glia for four weeks were positive for pre- and post-synaptic markers Synaptophysin I and PSD-95 (Figure S4A). Calcium imaging by FDSS 7000EX platform demonstrated spontaneous intracellular calcium oscillations, indicating that spontaneous firing was synchronized between the majority of the cortical neurons in the 96-wells (Figure S4B). Taken together these results suggest that the 3D spheroid protocol generate cortical neurons with expected transcriptional identity that continue to mature upon platedown with expression of synaptic markers and features of neuronal connectivity at network level [51].

Initial coarse-grained clustering using MCL identified four major groups of cells that specifically express known markers for NPs [52], mixed glial cells (G), CR cells [53] and neurons (N) [54] (Figure 6A,B). A small population of contaminating fibroblasts (0.1% of total cells) was removed from the dataset for downstream analyses. CR cells, expressed *DCX, CALB2, STMN2*, and *MAPT* consistently with developing mouse and human cortex (Figure 6B) [55–57]. The robust expression of *FOXG1* in the general population (Figure S5A) and the expression of *PAX6, EMX2*, and *LHX2* in NPs (Figure 6B) indicated our differentiation protocol mainly generates cells with dorsal telencephalic identity [58].

**Figure 6:**
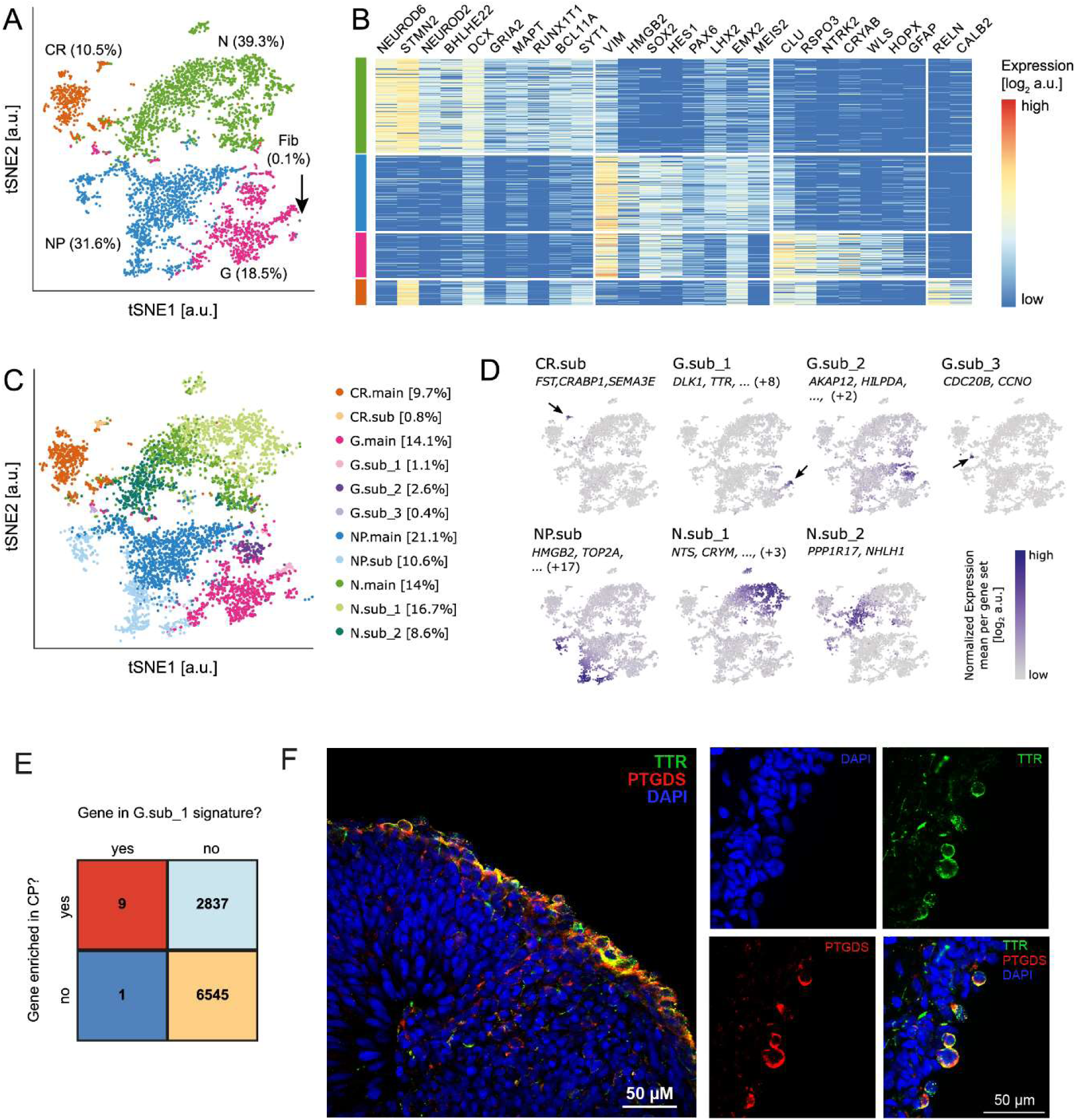
Characterization of hPSC derived cortical excitatory neurons by scRNA-seq. A: tSNE projection of 4857 single-cell trancriptomes of hPSC derived neuronal cell types after 86 days of differentiation. Unsupervised clustering using MCL groups cells into four major classes: Neurons (N), neuroepithelial progenitors (NP), mixed glial cells (G) and Cajal-Retzius cells (CR). In addition, a small population of fibroblasts (Fib) is identified. B: The identified cell populations are characterized by expression of known markers for the expected cell types. Expression values are shown as log2 (normalized UMI counts + 1). C: tSNE projection, colored by CellSIUS assignment. Main clusters are denoted .main, subclusters .sub. D: Mean expression of each marker gene set identified by CellSIUS, projected onto the same tSNE map as shown in A. The top markers are indicated for each gene sets, numbers in brackets refer to how many additional genes are part of the marker gene set. E: Comparison of the gene signature uncovered by CelSIUS to genes found to be enriched (p<0.05) in choroid plexus of the Fourth ventricle according to harmonizome[59,60]F: Single optical sections of neurosphere cryosections acquired by confocal microscopy showing co-localisation of TTR and PTGDS in cells predominantly on the periphery of neurospheres (panel left – compositie image of a neurosphere; panels right - split images from a different neurosphere).

Applying CellSIUS to this data identified 7 subpopulations (Figure 6C). Notably, within the mixed glial cells (G), CellSIUS identified a rare subgroup (1.1% of total population, G.sub_1) characterized by a signature of 10 genes. Nine of those ((TRPM3, PTGDS, TTR, CXCL14, HTR2C, WIF1, IGFBP7, MT1E, DLK1) are enriched in primary pre-natal human choroid plexus (CP) (Figure 6E) compared to the other tissues from the developing human cortex as defined in the harmonizome database [59,60] using a cutoff of 1.3 for the standardized value, sorresponding to p<0.05. This G.sub1 population is therefore consistent with formation of CP, a secretory neuroepithelial tissue that produces cerebrospinal fluid (CSF), and that has multiple origins along the rostro-caudal axis of the developing nervous system including the dorsal telencephalic midline [61]. We further validated the presence of CP neuroepithelia in our 3D human cortical cultures by confocal microscopy analysis. Using neurosphere cryosections, we demonstrated co-localisation of canonical CP marker Transthyretin (TTR) with Prostaglandin D2 Synthase (PTGDS), another CP enriched protein described in primary mouse and human tissue, in a limited number of cells located almost exclusively on the periphery of neurospheres (Figure 6F). Collectively these results suggest that the 3D spheroid human cortical differentiation protocol described here can generate developmentally relevant cell types and that CellSIUS can identify rare cell populations within the heterogeneity and complexity of stem cell-based models. CellSIUS identified a second subgroup in the mixed glial cells (G) characterized by high expression levels of glycolytic enzymes (G.sub_2, 2.6%) (Figures 6C,D and S6A). analysis between G.sub_2 and the rest of the G cells revealed upregulation of *HOPX, PTPRZ1, CLU, BCAN, ID4*, and *TTYH1* in the main group, a transcriptional signature consistent with developing human outer radial glia (oRG) [62], (Table S4, Figure S6A). oRG cells also upregulated mitochondrial genes (Table S4) that are crucial for oxidative phosphorylation, highlighting the metabolic difference between these two groups. We hypothesize the G.sub_2 subgroup to be a progenitor population that is located closer to the hypoxic interior of neurospheres, a common feature of the 3D spheroid differentiation protocols.

In addition, CellSIUS identified a subgroup of NP cells (NP.sub, 10.6%) defined by upregulation of cell-cycle related genes such as *HMGB2, TOP2A* and *MKI67* (Figures 6C,D and S6A) as well as a subgroup of CR cells (CR.sub, 0.8%) characterized by *SEMA3E, BTG1*, and *PCDH11X* (Figures 4B and S5) which may represent CR cells at a different stage of migration [63–65]. Finally, CellSIUS revealed a split in the neuronal population (N), identifying 2 groups, N.sub_2 (8.6%) and N.sub_1 (16.7%) (Figures 6C, D and S6A). In addition to *NHLH1* and *PPP1R17* known to be enriched in immature neurons [62], N.sub_2 expressed *EOMES* (Figure S5B), a well characterized marker of cortical intermediate progenitors [54,66] that give rise to TBR1+ cortical neurons (Figure S5C) and is likely a mixed population of intermediate progenitors and immature neurons. In contrast, markers identified by CellSIUS for the N_sub1 neuronal population were unexpected. Although co-expression of *FEZF2, CRYM, PCDH17* and *RUNX1T1* in this cortical neuronal population is consistent with recent scRNA-seq data from the developing human cortex (Figure S7B, EN-V1-1: Early-born deep-layer/sub-plate excitatory neurons, EN-PFC1: Early-born deep-layer/sub-plate excitatory neurons prefrontal cortex), robust *NTS* expression in developing cortical neurons has not been reported so far to the best of our knowledge. The expression of FEZF2 (Figure S5D) in this culture is consistent with the general dorsal telencephalic identity of these cells and co-expression of FEZF2 and BCL11B (CTIP2) in this particular post-mitotic neuronal sub-population (Figure S5E) could suggest patterning towards cortico-spinal motor neurons (CSMNs). However, the presence of NTS, which encodes a 13 amino acid neuropeptide called neurotensin highly expressed in the hypothalamus and amygdala, is not in line with the overall transcriptional identity as discussed above. Analysis of a recently published scRNA-seq dataset from different regions and developmental stages of the human cortex [54] revealed that only a few cells derived from the fetal primary visual cortex (age 13 pcw) express *NTS* (Figure S7B). However, the number of cells in our dataset were too low to draw any firm conclusions.

To further characterize the transition from progenitors to the two different neuronal cell types (CR cells and all N populations), we applied Monocle for trajectory analysis to a subset of the cells corresponding to these three identities. This analysis revealed a tree with two branches (Figure 7A). As expected, cells progress from the tree root which is composed of progenitors *via* the NHLH1^high^/PPP1R17^high^ population towards either N (branch 1) or CR cells (branch 2). Along the trajectory, the NP marker *VIM* decreases gradually whereas *NHLH1* increases up to the branch point, then decreases again (Figure 7B). The CR branch ends with cells expressing high levels of *RELN*, and the N branch is characterized by gradual increase of *FEZF2* expression and ending in the N.sub_1 population (Figure 7B). Notably, at the very tip of this branch, we also find a very small number of cells expressing *LDB2* and *DIAPH3* which are markers of CSMNs in the mouse [67]. It is plausible that, given more time, this population may eventually give rise to CSMNs with a more defined transcriptional signature.

**Figure 7:**
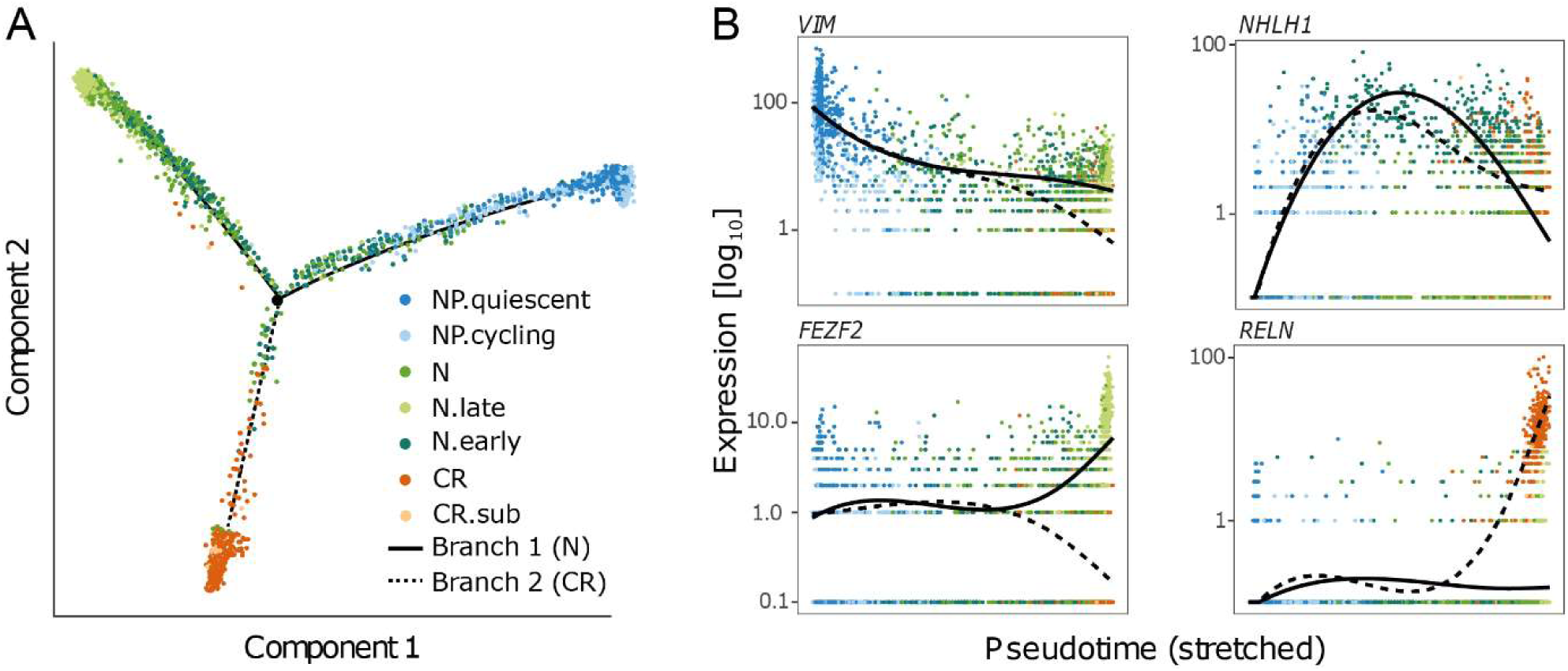
Monocle analysis of the NP, N and CR cluster. A: Consistent with the subgroup assignment by CellSIUS, monocle orders cells on a trajectory from NP via immature neurons (N_early) to either mature N or CR cells. B: Gene expression along pseudotime. Shown are a marker for NPs (VIM), immature neurons (NHLH1), N.sub_2 (FEZF2) and CR cells (RELN).

### Comparison of CellSIUS, RaceID and Giniclust2 performance for rare cell type identification in hPSC-derived cortical neurons

Finally, to compare CellSIUS’ performance for rare cell type identification in complex and heterogenous stem cell data, we compared its output to GiniClust2 and RaceID3 results. Application of GiniClust2 to the hPSC-derived cortical neurons initially grouped by MCL into 4 main clusters resulted in a total of 20 clusters. The main differences between GiniClust2 and CellSIUS (Figure S6B) results can be summarized as follow: GiniClust2 generated clusters that merge major known cell types (for example cluster 24 merges glia, glia_1 (=CP), glia_2, neurons, N.sub-1 (late neurons) and N.sub_2 (early neurons)), (ii) GiniClust2 did not detect CP (G.Sub_1), cycling NPs (NP.sub) nor the well described immature neurons (N.sub_2). Application of RaceID to the hPSC-derived cortical neurons initially grouped by MCL into 4 main clusters resulted in a total of >50 clusters with default parameters consistently with the high false positive rate observed with synthetic and cell line data. With a more stringent outlier probability cutoff (10^-20), RaceID3 identifies 10 clusters with a similar overall assignment to CellSIUS (Figure S6C). However, if RaceID3 did detect CP (G.Sub_1), it split this cluster across several other clusters with the majority of cells assigned to either cluster 3 (19 CP together with 4 other cells) or cluster 5 (mixed with a large number of G, N and NP cells). The CP markers *PTGDS* and *TTR* are co-expressed in 49/53 CP cells identified by CellSIUS but only in 19/54 CP cells identified by RaceID3 suggesting that RaceID3 incorrectly assigned most of the CP cells to a merged glia / NP / N cluster. In addition, and similarly to GiniClust2, RaceID3 did neither identify cycling NPs (NP.sub) nor the above described progenitors and immature neurons population (N.sub_2).

In summary, we show that CellSIUS demonstrates superior performance for specificity and sensitivity compared to other approaches in complex and heterogenous data and enables the identification of rare populations as small as 0.4% within major cell types that differ by their metabolic state, cell cycle phase, or migratory state.

## Discussion

We generated a comprehensive benchmark dataset of ∼12 000 single cell transcriptomes from 8 cell lines to evaluate the performance of scRNA-seq feature reduction and clustering approaches. Our findings suggest that for unsupervised feature selection, the DANB methods implemented in the M3Drop package outperformed HVG. Whilst all clustering methods tested performed equally well on data with balanced and abundant cell populations, k-means and model-based methods performed poorly on subsampled datasets with unequal cell type proportions, typically splitting clusters containing many cells while merging those containing few cells. This is likely a consequence of feature selection and PCA-based dimensionality reduction prior to clustering where these methods select or assign weights to genes based on mean expression and variance across the whole cell population, which are both low if a gene is specifically expressed in a small subset of cells only.

In contrast, hclust in combination with dynamicTreeCut, MCL and DBSCAN resulted in accurate cluster assignments across all subsampled datasets. Strikingly, none of the methods we tested was able to identify rare cell types (<1%). It is worth noting that although DBSCAN does classify rare cell types as border points, it does not reliably identify these populations for two reasons: (i) additional cells that did not belong to the rare populations are also classified as border points; (ii) DBSCAN does not perform well if there are points connecting clusters, which is often the case in scRNA-seq datasets.

To overcome these limitations, we developed CellSIUS, a novel algorithm that takes initial coarse clusters as input and identifies rare cell subtypes based on correlated gene sets specific to subpopulations. The overall idea behind CellSIUS is similar to RaceID3 [46] and GiniClust2 [20], two recent methods for the identification of rare cell types in scRNA-seq datasets. All of these algorithms combine a global clustering with a second assignment method which is tailored to finding rare cell types. There are however, important differences between the approaches which are at the basis of CellSIUS’ superior performance for both rare cell type as well as outlier genes identification in terms of specificity and selectivity.

RaceID3’s initial step is a k-medoids clustering, followed by outlier cell identification in each cluster in four steps: (i) calibration of a background model of gene expression by fitting a negative binomial distribution to the mean and variance of each gene in each cluster; (ii) identification of outlier cells by calculating for each gene and each cell the probability of observing this expression value under the assumption of the background model; (iii) merging of potential outlier cells into new clusters based on the similarity of their gene expression; and (iv) definition of new cluster centers for both the original and the outlier clusters. In a final step, cells are assigned to the cluster they are closest to. In contrast to CellSIUS, RaceID3 does not require the outlier genes to be cluster specific; consequently, it may select genes that co-vary with technical confounders such as the total number of detected genes per cell. In addition, whereas CellSIUS only considers subcluster-specific genes to assign cells to final clusters, the final cluster assignment in RaceID3 is done based on the similarity of each cell’s whole transcriptomic signature to each cluster center. In cases where the distance between the outlier cluster and neighboring clusters is small, this leads to a high number of false positives, with many cells initially not identified as outliers being merged into the nearest outlier cluster.

GiniClust2 runs two independent clustering steps on the same data. The first clustering aims at capturing global structure of the data by running a k-means clustering on the expression of genes with a high Fano factor. This is motivated by the fact that a high Fano factor is associated with genes that are differentially expressed between abundant cell types. The second clustering is performed by running a density based clustering on genes with a high Gini index which is typically associated with genes being differentially expressed between rare and abundant cells. In a final step, the results of both clustering are merged based on a weighted consensus association. The main differences to CellSIUS are as follows: (i) the selection of the genes for the rare cell type assignment is performed using a global metric (i.e. the Gini coefficient across the whole dataset), whereas CellSIUS takes into account the information on the global clustering (e.g. considers only cluster specific genes); (ii) the final assignment is a weighted average of the results from both clustering steps, whereas we use a two-step approach consisting of an initial coarse clustering step followed by CellSIUS for the identification of rare cell types and outlier genes.

In addition to CellSIUS’ superior performance described above, which potentially reflects the propensity of RaceID3 and GiniClust2 to interpret technical variation as biological signal in single cell transcriptomic data, our novel approach simultaneously reveals transcriptomic signatures indicative of rare cell type’s function.

In order to use our methods in a real-world setting, we applied the workflow presented here to a dataset from hPSC derived neurons and identified major neural cell types of early human corticogenesis such as cycling and quiescent NPs, *EOMES*^+^ IPs, CR cells, immature and mature neurons with a transcriptional identity indicative of layer V/VI neurons, and oRG. Overall, the transcriptional fingerprint of each major group was in line with a recent scRNA-seq data set from the developing human cortex. CellSIUS analysis also revealed a transcriptional signature in the mature neuronal population that begins to deviate from the expected cortical trajectory, typified by the high expression levels of *NTS* detected in N.sub_1, highlighting the importance of unbiased characterization of hPSC differentiation platforms at single cell level. Single-cell trajectory analysis of NP, CR and N cells using Monocle revealed a pseudo-temporal order of progenitors gradually differentiating into neurons, with a lineage split between Cajal-Retzius cells and *FEZF2*^+^ neurons.

Importantly, CellSIUS analysis identified rare cell types within the major groups, such as putative CP (G.sub_1) making up 1.1% of the cell population, which were not identified by existing approaches for rare cell type identification. We validated the presence of CP neuroepithelia in our 3D cortical spheroid cultures by confocal microscopy and cross-referenced CP-specific gene list identified by CellSIUS to primary pre-natal human data. In addition, CellSIUS analysis provided a signature gene list for human PSC-derived CP cells *in vitro* for the first time, paving the way for isolation, propagation, and functional characterization of this lineage.

One drawback of CellSIUS is that it is sensitive to the initial cluster assignments. In practice, this should only be an issue if there is no clear global structure in the data and cluster assignments are not consistent between different clustering methods and/or parameter settings. In such cases, one could use a consensus assignment from a combination of different clustering assignments. In summary, we developed, benchmarked and implemented CellSIUS, a novel method for detection and characterization of rare cell types from complex scRNA-seq data. The large single-cell RNA-seq dataset of known cell composition generated for this work, represents a biological ground truth for benchmarking of future novel methods. We exemplify the use of CellSIUS for the characterization of a novel human pluripotent cell differentiation protocol recapitulating deep-layer corticogenesis *in vitro*. scRNA-seq in combination with highly sensitive and specific computational approaches such as CellSIUS provide an unprecedented resolution in the transcriptional analysis of developmental trajectories, revealing previously unrecognized complexities in human stem cell-derived cellular populations. This study represents a rich dataset as benchmark for derivation of cortical neurons from human PSCs using small molecules, can inform refinement of directed-differentiation approaches to ultimately generate *bona fide* CSMNs and upper-layer excitatory neurons, and enable isolation and characterization of CP neuroepithelia that are crucial to study neurological disorders *in vitro*.

## Methods

### Human cell lines

For the benchmarking dataset, 8 different human cell lines from the ATCC biorepository have been used (Table 2).

### Single-cell RNA-sequencing of cell lines

Cellular suspensions were loaded on a 10x Genomics Chromium Single Cell instrument to generate GEMs. Single-cell RNA-seq libraries were prepared using GemCode Single Cell 3’ Gel Bead and Library Kit according to CG00052_SingleCell3’ReagentKitv2UserGuide_RevB. GEM-RT was performed in a Bio-Rad PTC-200 Thermal Cycler with semi-skirted 96-Well Plate (Eppendorf, P/N 0030 128.605): 53 °C for 45 minutes, 85 °C for 5 minutes; held at 4 °C. After RT, GEMs were broken and the single strand cDNA was cleaned up with DynaBeads^®^ MyOne™ Silane Beads (Life Technologies P/N, 37002D). cDNA was amplified using a Bio-Rad PTC-200 Thermal cycler with 0.2ml 8-strip non-Flex PCR tubes, with flat Caps (STARLAB, P/N I1402-3700): 98 °C for 3 min; cycled 12x: 98 °C for 15 s, 67 °C for 20 s, and 72 °C for 1 min; 72 °C for 1 min; held at 4 °C. Amplified cDNA product was cleaned up with the SPRIselect Reagent Kit (0.6X SPRI). Indexed sequencing libraries were constructed using the reagents in the Chromium Single Cell 3’ library kit V2 (10x Genomics P/N-120237), following these steps: 1) Fragmentation, End Repair and A-Tailing; 2) Post Fragmentation, End Repair & A-Tailing Double Sided Size Selection with SPRIselect Reagent Kit (0.6X SPRI and 0.8X SPRI); 3) adaptor ligation; 4) post-ligation cleanups with SPRIselect (0.8X SPRI); 5) sample index PCR using the Chromium Multiplex kit (10x Genomics P/N-120262); 6) Post Sample Index Double Sided Size Selection-with SPRIselect Reagent Kit (0.6X SPRI and 0.8X SPRI). The barcode sequencing libraries were quantified using a Qubit 2.0 with a Qubit ™ dsDNA HS Assay Kit (Invitrogen P/N Q32854) and the quality of the libraries were performed on a 2100 Bioanalyzer from Agilent using an Agilent High Sensitivity DNA kit (Agilent P/N 5067-4626). Sequencing libraries were loaded at 10pM on an Illumina HiSeq2500 with 2 × 50 paired-end kits using the following read length: 26 cycles Read1, 8 cycles i7 Index and 98 cycles Read2. The CellRanger suite (2.0.2) was used to generate the aggregated gene expression matrix from the BCL files generated by the sequencer based on the hg38 Cell Ranger human genome annotation files.

### Bulk RNA-sequencing of cell lines

For each individual cell line, RNA was isolated from 5×10^5^ cells using the RNeasy Micro kit (Qiagen, Cat# 74104). The amount of RNA was quantified with the Agilent RNA 6000 Nano Kit (Agilent Technologies, Cat# 5067-1511). RNA sequencing libraries were prepared using the Illumina TruSeq RNA Sample Prep kit v2 and sequenced using the Illumina HiSeq2500 platform. Samples were sequenced to a length of 2×76 base-pairs. Read pairs were mapped to the Homo sapiens genome (GRCh38) and the human gene transcripts from Ensembl version 87 [68] by using an in-house gene quantification pipeline [69]. Genome and transcript alignments were used to calculate gene counts based on Ensembl gene IDs.

### Differentiation of cortical excitatory neurons from human pluripotent stem cells in suspension

H9-hESCs (WA09) were obtained from WiCell and maintained in TeSR-E8 medium (Stemcell Tech., 05990) on tissue-culture plates coated with vitronectin (Gibco, A14700). hESCs were passaged using ReLeSR (Stemcell Tech., 05873) to dissociate into cell clumps and were replated in E8 plus thiazovivin (Selleckchem, S1459) at 0.2μM. H9-hESC line was free of myoplasma and was tested using the Mycoalert detection kit (Lonza).

hESCs were changed to mTesR1 (Stemcell Tech., 85850) media when they were 70-80% confluent and maintained in mTesR1 for minimum of two days before confluent monolayer of hESCs were neurally converted by changing the media to Phase I (Table S5). Seven days post induction, cells were dissociated to single-cell suspension with Accutase (Gibco A1110501), seeded at 1.5E6 cells /mL in spinner flasks with Phase II media (Table S5) supplemented with 2 μM Thiazovivin and 10 ng/mL FGF2 (Peprotech, 100-18B) (final) and incubated at 37°C on a micro-stir plate at 40 rpm for 4 days. Media was then changed to Phase III (Table S5) and neurospheres were further cultured for 17 days at 60 rpm, changing media 50% twice a week. On day 28 media was changed to Phase IV (Table S5) and cultures were maintained 21 more days with 50% media change twice a week. From day 49 onwards cultures were switched to Ph IV media for maintenance. Neurospheres were dissociated with Papain kit (Worthington) at day 86 for single-cell RNAseq or neuronal platedowns on laminin (Sigma, L2020), fibronectin (Corning, 354008), and Matrigel (Corning, 354230) coated plates.

### Immunofluorescence and cryosectioning

Cells were fixed with 4% PFA, permeabilised with 0.2% Triton X-100 at room temperature, and then blocked in 3% goat serum, followed by incubation with primary (TBR1 - Abcam, ab31940; CTIP2 – Abcam, ab18465; β3 tubulin – Biolegend, 801202; PSD-95 – Synaptic Systems, 124 011; Synaptophysin 1 – Synaptic Systems, 101 002; Transthyretin – Novus Biologicals, NBP2-52575, Prostaglandin D Synthase (PTGDS) – Abcam, ab182141) and secondary antibodies (Alexa Flours, Invitrogen). The nuclei were counter-stained with 49,6-diamidino-2-phenylindole (DAPI, Sigma). Cryosectioning of neurospheres were performed as previously described [70]. Cells were imaged using an Observer D1 (Zeiss) microscope or Olympus SD-OSR spinning-disk confocal microscope (60x oil immersion). The images were processed using Zen 2 (Zeiss), MetaMorph or Image J (brightness and contrast adjustments, thresholding for composite images) and assembled using Adobe Photoshop CS6.

### Calcium imaging

The intracellular Ca^2+^ oscillations in human cortical neuron and rat glia co-cultures were assessed using the FLIPR Calcium 6 Kit (Molecular Devices LLC, San Jose, California). Briefly, 96-well Greiner μ-clear plates (655097) were seeded with 2500 rat glia (Lonza, R-CXAS-520) per well in Ph IV media and cultured for seven days. Human cortical neurospheres were dissociated with papain as described above at DIV 56 and 50,000 single cells per well were plated on rat glia in Phase IV media. Co-cultures were maintained for four weeks with twice weekly 50% media exchange. Cells were loaded with Calcium 6 dye for an hour which was reconstituted in imaging buffer (NaCl 2.5 mM, KCl 125 mM, KH_2_PO_4_ 1.25 mM, CaCl_2_ 2 mM, MgCl_2_ 2 mM, HEPES (acid) 25 mM, D-glucose 30 mM, pH 7.4, filter-sterilised). Kinetics of Ca^2+^ oscillations were determined as fluorescence intensity at 540 nm following excitation at 480 using the FDSS 7000EX Functional Drug Screening System (Hamamatsu) maintained at a constant 37°C throughout the assay. A total of 3000 reads per assay were recorded. The exposure time per read was 100 ms with sensitivity set to 1.

### Single-cell RNA-sequencing of neuronal cells

Cells were resuspended to 1 million cells/mL and run through the 10X Chromium, Version 2 single cell RNA-seq pipeline per vendor’s instructions. Reverse transcription master mix was prepared from 50µL RT reagent mix (10X, 220089), 3.8µL RT primer (10X, 310354), 2.4µL additive A (10X, 220074), and 10µL RT enzyme mix (10X, 220079). 4.3µL cell solution was mixed with 29.5µL H2O and 66.2µL reverse transcription master mix. 90µL sample was loaded onto the 10X Single Cell 3’ Chip along with 40µL barcoded gel beads and 270µL partitioning oil, and the microfluidics system was run to match gel beads with individual cells. The droplet solution was then slowly transferred to an 8-tube strip, which was immediately incubated for 45 minutes at 53°C to perform reverse transcription, then 5 minutes at 85°C. The sample was treated with 125µL recovery agent (10X, 220016), which was then removed along with the partitioning oil. 200µL of cleanup solution containing 4µL DynaBeads MyOne Silane Beads (Thermo Fisher, 37002D), 9µL water, 182µL Buffer Sample Clean Up 1 (10X, 220020), and Additive A (10X, 220074) was added to the sample, and the solution was mixed 5 times by pipetting and allowed to incubate at room temperature for 10 minutes. Beads were separated via magnetic separator and supernatant was removed. While still on the magnetic separator, the beads were then washed twice with 80% ethanol. The separator was then removed and the beads were resuspended in 35.5µL elution solution consisting of 98µL Buffer EB (Qiagen, 19086), 1µL 10% Tween 20 (Bio-Rad, 1610781), and 1µL Additive A (10X, 220074). The solution was then incubated for 1 minute at room temperature, and placed back onto the magnetic separator. 35µL of eluted sample was transferred to a new tube strip. cDNA amplification reaction mix was prepared from 8µL water, 50µL Amplification Master Mix (10X, 220125), 5µL cDNA Additive (10X, 220067), and 2µL cDNA Primer Mix (10X, 220106). 65µL of amplification master mix was added to the sample, mixed 15 times via pipetting, and briefly centrifuged. The sample then underwent 12 amplification cycles (15 seconds at 98°C, 20 seconds at 67°C, 1 minute at 72°C).

SPRIselect beads (Beckman Coulter, B23318) were then applied at 0.6X, and solution was mixed 15 times via pipetting. The sample was incubated at room temperature for 5 minutes, placed onto a magnetic separator, and washed twice with 80% ethanol. Sample was air dried for 2 minutes and eluted in 40.5µL Buffer EB. cDNA yield was measured on a 2100 Bioanalyzer (Agilent, G2943CA) via DNA High Sensitivity Chip (Agilent, 5067-4626).

Fragmentation mix was prepared at 4°C from 10µL fragmentation enzyme blend (10X, 220107) and 5µL fragmentation buffer (10X, 220108). 35µL of sample cDNA was then added to the chilled fragmentation mix. Sample was incubated for 5 minutes at 32°C, then 30 minutes at 65°C to conduct enzymatic fragmentation, end repair, and A-tailing. Sample was then purified using 0.6X SPRIselect reagent (see above). Adaptor ligation mix was prepared from 17.5µL water, 20µL Ligation Buffer (10X, 220109), 10µL DNA Ligase (10X, 220110), and 2.5µL Adaptor Mix (10X, 220026). The ligation mix was added to 50µL of sample and mixed 15 times via pipetting. Sample was then incubated for 15 minutes at 20°C to conduct the ligation. The sample was purified using 0.8X SPRIselect reagent (see above). Sample index PCR mix was prepared from 8µL water, 50µL Amplification Master Mix (10X, 220125), and 2µL SI-PCR Primer (10X, 220111). 60µL sample index PCR mix, 30µL purified sample, and 10µL of sample index (10X, 220103) were combined and mixed 15 times via pipetting. Indexing was conducted via 9 cycles of 20 seconds at 98°C, 30 seconds at 54°C, then 20 seconds at 72°C. Sample was purified via double-sided SPRI selection at 0.6X and 0.8X, respectively. Sample was then quantified via DNA High Sensitivity Chip.

Additional quantification was conducted via KAPA Library Quantification Kit (Illumina, KK4828- 07960166001). Sample was diluted at 10-fold increments from 1:100 to 1:1,000,000, and mixed 1:9 with KAPA qPCR mix. qPCR was conducted on a Viia7 qPCR machine (Life Technologies).

Sample was then sequenced on a HiSeq 4000 (Illumina) using 2 x 50-cycle SBS kits (Illumina, FC-410- 1001). Sample library was diluted to 2nM in EB buffer with 1% PhiX spike-in. 5µL nondenatured library was then mixed with 5µL 0.1N NaOH, then vortexed and briefly centrifuged. Denaturing was conducted at room temperature for exactly 8 minutes, then stopped via addition of 5µL 200mM Tris-HCl pH 8.0 (Fluka, 93283). Sample was mixed, briefly centrifuged, and placed on ice. ExAmp reaction mix (Illumina, PE-410-1001) was prepared, added to the sample, and clustering was done on a HiSeq 4000 flow cell via cBot2 (Illumina). The library was then sequenced with paired-end reagents, with 26xRead 1 cycles, 8xi7 index cycles, and 98xRead 2 cycles.

The 10X Cell Ranger 1.3.1 pipeline was utilized to convert raw BCL files to cell-gene matrices. FASTQ files were aligned to the GRCh37.75 human reference genome, UMI-filtered, and barcodes were matched via the CellRanger count script

### Computational analysis

#### Software requirements

All computational analysis was carried out using R v. 3.4.1 with Bioconductor v. 3.5.

### Generation of synthetic data

A synthetic dataset was generated based on estimated parameters for the gene-wise mean μ_*i*_ and variance 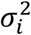 from experimentally determined counts of 1000 K562 cells from our benchmarking dataset.

Because gene expression within each cell is typically not independent but cells that have high/low count number for one gene also tend to have high/low counts for another, we sampled for each cell j a scaling factor *θ*_*j*_ such that log_2_(*θ*_*j*_)∼ 𝒩 (0,0.25), as described in [31]. Simulated counts for gene *i* and cell *j* were generated by sampling from a negative binomial with mean

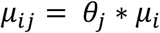

and dispersion^1^

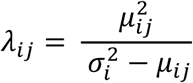

A second order polynomial was fit to the sample variance as a function of the mean in logarithmic space as described in [9]. This polynomial served as an estimate of the global mean-variance relationship. Replacing the term 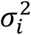 in the equation above with this estimate, the dispersion can be expressed as a function of *μ*_*ij*_:

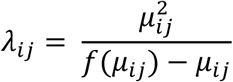

Where

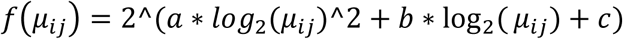

is derived from the second order polynomial approximating the gene-wise variance as a function of mean expression. For genes exhibiting Poissonian behavior (i.e. equal mean and variance), we set *λ* to a fixed value of 10^10^.

Main cell populations were obtained by permutation of the expression values of 100 randomly chosen genes with mean counts larger than 2.

Cell subgroups characterized by high expression of a small set of marker genes were generated by replacing the base mean values μ_*i*_ in a small set of genes with low expression (μ_*i*_ < 0.1) by a value of 2^*x*^ where *x* ∼ *𝒩*(2.5,1). Thus, the upregulated genes exhibit a log2 fold change of 2.5 on average.

### Simulating varying degrees of subtlety in transcriptional differences

An initial small dataset was subsampled from the benchmarking (8 human cell lines) dataset, comprising 100 HEK293, 125 Ramos, and between 10 Jurkat cells. We used scran to predict cell sycle stage and only included cells in G1 phase.

From this initial dataset, 25 Ramos cells were held out. From the remaining dataset (100 HEK293, 100 Ramos, 10 Jurkat), datasets with varying incidence of a rare cell type and subtlety of its transcriptional signature were generated in silico, following an approach recently described by Crow et. al[47]: First, a number of Jurkat cells (i.e. incidence of 2,5 or 10) were sampled form the initial dataset. Then, to simulate varying degrees of transcriptional difference between the rare cell type (Jurkat) and its closest abundant cell type (Ramos), an increasing fraction of gene expression values, ranging form 0 to 0.995 in steps of 0.05 (0.045 for the very last step) in the Jurkat cells were replaced by the respective values in the held out Ramos cells.

This procedure was repeated 5 times for each incidence of the rare cell type and each value of the subtlety parameter.

The performance of CellSIUS, GiniClust2 and RaceID3 was evaluated in terms of recall, precision and true negative rate (TNR) for each configuration. To this end, a confusion matrix between the true cell type and the predicted cell type was generated. “Main clusters” were defined as the two clusters containing the majority of the HEK293 and Ramos cells, respectively. The TPR was then defined as the fraction of Jurkat cells that were not assigned to the main clusters, precision was defined as the fraction of Jurkat cells among all cells not assigned to the two main clusters, and the TNR was defined as the fraction of HEK293 and Ramos cells that were assigned to the main clusters.

### Data pre-processing

Initial pre-processing was applied to each batch of cell lines separately prior to annotating cell types.

First, cells were filtered based on the total number of detected genes, total UMI counts and the percentage of total UMI counts attributed to mitochondrial genes. Cutoffs were set individually per batch based on the overall distributions (Table S3).

Second, genes have to present with at least 3 UMIs in at least one cell. After this initial QC, remaining outlier cells were identified and removed using the *plotPCA* function from the scater [30] R package with *detect_outliers* set to TRUE.

Data were normalized using scran [31], including a first clustering step as implemented in the *quickCluster* function and with all parameters set to their default values.

### Cell type annotation

First, the top 10% overdispersed genes were selected using the NBDrop method described in [37]. Cell types were then annotated based on Pearson correlation of the expression profile (log_2_(normalized counts+1)) of the selected features with bulk RNA-seq data obtained for each individual cell line (Figure 1A-B). For the batches 1-3 that contained only two cell lines each, the Pearson correlation coefficients were scaled to z-scores prior to the assignment, for batch 4, the raw correlation values were used instead. A cell was then assigned to the cell line with the highest value unless this maximum was below 0.2 or if the second highest value was within 5% of the maximum in which case no assignment was given. We found that the latter applied only to a small percentage of cells (1–2%), which most likely correspond to cell doublets. Furthermore, for the cell line mixes IMR90/HCT116 and A549/Ramos additional potential doublets were identified and excluded from the cell line assignment employing a visual inspection of the tSNE plot by looking for (small) clusters of cells having high correlation to both cell lines as well as a high UMI count (Table S3).

After cell type annotation, the raw count matrices from all four batches were concatenated. Cells that had not passed the initial QC or could not be annotated were discarded. The gene filtering step described above was then repeated for the aggregated dataset, leaving a final cleaned dataset containing a total of 12 718 genes and 11 678 cells.

### Dimensionality reduction and calculation of distance matrix

The original expression (log2(normalized counts + 1) coordinates were projected into low dimensional space by PCA, using an implicitly restarted Lanczos method as implemented in the irlba [44] R package. The number of dimensions to retain was determined by visual inspection of a screeplot. It was 10 for all cell line data and 12 for the neuron dataset, and the first k principal components accounted for 40–50% of the total variance in each case. Cell-cell distances (Euclidean or Pearson, Table 2) were then calculated on these projections.

### Benchmarking of clustering approaches

The accuracy of each prediction was assessed by the adjusted rand index (ARI). Given two partitions *X* = *X*_1_, …, *X*_*m*_ and *Y* = *Y*_1_, …, *Y*_*k*_ of a set S with *n* elements, the ARI is defined as

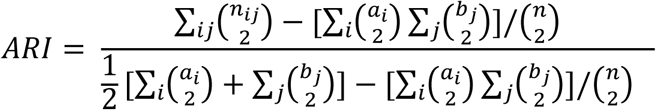

Where *n*_*ij*_ denotes the elements that are common between *X*_*i*_ and *Y*_*j*_, and *a*_*i*_, *b*_*j*_ are the total number of elements in *X*_*i*_ and *Y*_*j*_, respectively.

### CellSIUS

CellSIUS detects cell subpopulations and their gene signatures (Figure 3A). Starting from an initial partitioning of *N* cells into *m* clusters *C*_1_, …, *C*_*m*_, the method identifies cell subpopulations and their signatures as follows:

1. Identification of genes with bimodal expression: For each gene *g*_*n*_, within each cluster *C*_7_, a 1-dimensional k-means clustering is used to partition the cellular expression levels (log2 normalized UMI counts) into two groups (“low” and “high”). Candidate marker genes are selected according to three criteria: (i) the average expression fold change between “low” and “high” is at least 2 on a log2-scale, (ii) less than a user defined percentage (50% by default) of all cells in cluster *C*_*j*_ fall in the “high” category, (iii) there is a significant difference (t-test and Benjamini-Hochberg correction, p-value < 0.1) between the “low” and “high” expression values.
2. Testing cluster specificity: For the list of candidate genes, it is assessed whether the cell subgroup expressing them is specific to cluster *C*_*j*_. Required for each gene *g*_*i*_ are (i) a significant difference in the expression of *g*_*i*_ in cells with “high” expression compared to cells not in Cj (t-test and FDR correction, p-value < 0.1), and (ii) the average expression fold change between all cells with “high” expression and all other cells with non-zero expression of *g*_*i*_ to be at least 1 on a log2-scale.
3. Identification of correlated gene sets: For each cluster *C*_*j*_, the correlation matrix of the expression of all candidate genes *g*_1,..,*n*_ across all cells in cluster *C*_*j*_ is transformed into a graph where genes correspond to nodes and edges are weighted by correlations between them. Edges with weights below a fixed threshold are assigned a weight of 0. By default, this threshold is set to the 95^th^ percentile of all correlations if this value lies between 0.35 and 0.5, and to the lower and upper bound if it is below or above, respectively. The lower bound is set such that it is higher than the maximum of all gene-wise correlations on simulated data from an entirely homogeneous population, which serves as an estimate of the background correlation. Setting an upper bound ensures that gene sets are not falsely split in cases where all candidate genes are highly correlated. Subsequently, MCL [41,42] is used to identify correlated gene sets, denoted *s*_*jk*_, where *j* is the index of the main cluster and *k* the index of the gene set within this cluster.
4. Assigning cells to subgroups: For each cluster *C*_*j*_ and each gene set *S*_*jk*_, a 1-dimensional k-means is run on the mean expression of *s*_*jk*_. Cells falling in the “high” mode of this clustering are assigned to a new cluster *C*_*jk*_.
5. Final cluster assignment: Cells are assigned to a final cluster which is the combination of all subgroups they belong to. Only subgroups characterized by a minimum of min_n_genes (default: 3 genes) are considered.

### Identification of rare cell types with RaceID and Giniclust

RaceID3 [46] was obtained from github (dgrun/RaceID3_StemID2, version as of March 26^th^ 2018). Analysis was run with all parameters at their default values, except that we fixed the initial clusters (RaceID@kpart) instead of determining them by k-medoids. On biological data (cell line subset 2 and neuronal population), we in addition changed the probability threshold to 10^-20^ and set the minimum number of outlier genes (outlg) to 3.

GiniClust2 [20] was obtained from github (dtsoucas/GiniClust2, version as of 4^th^ May 2018). All analysis was run with dataset specific parameters: MinPts = 3, eps = 0.45, k=2 for the simulated data and MinPts = 3, eps = 0.45, k=8 for the cell line dataset. All other parameters were set to their defaults.

### Trajectory analysis using Monocle

Analysis was run using monocle version 2.4.0. As input, the counts of the top 10% genes selected by NBDrop were used. Prior to monocle analysis, all genes annotated with the GO term cell cycle (GO:0007049) as well as mitochondrial genes and genes encoding ribosomal proteins were removed from the dataset. All parameters were set to default values.

## Code and Data availability

The code and processed data to reproduce the analyses presented here are included in this published article (see compressed supplementary folder). Raw data will be deposited to the NCBI Sequence Read Archive (SRA) upon publication. The workflow and CellSIUS are written in the R programming language. CellSIUS is provided as a standalone R package. It requires R >= 3.4.1 and uses an external installation of the Markov Clustering Algorithm (MCL) [41,42]. The R implementation is platform independent, the external MCL runs on any UNIX platform. The code, vignette and an example dataset for the computational workflow are included in this published article (see compressed workflow folder). The code and processed data will be available on github under the GNU GPL license upon publication).

**List of Abbreviations**
scRNA-seq: single-cell RNA sequencing
DE: differential expression
hPSC: human pluripotent stem cell
HVG: high variance gene
DANB: depth-adjusted negative binomial
PCA: principal component analysis
GMM: Gaussian mixture model
ARI: Adjusted Rand index
NP: neocortical progenitor
CR: Cajal-Retzius
IP: intermediate progenitor
N: neuron
oRG: outer radial glia
G: glia
CP: choroid plexus
GC: glycolytic cell

## Declarations

### Ethics approval

Not applicable.

### Consent for publication

Not applicable.

### Author contribution

MN, FN and RW developed CellSIUS and implemented the computational workflow. AW and RC sequenced the human cell lines for the benchmarking study. RW performed the benchmarking analysis. RW, MN, MS, BB and CGK analyzed and interpreted the neuroscience data. HN, MS and JR performed the experiments. MF, BB and AK designed experiments. MN, GR, SS, AJ, BB and CGK contributed to the conception of the studies and the interpretation of data. RW, MN, BB, MS, AK and CGK wrote the manuscript. All authors examined the results and approved the final version of the manuscript.

## Acknowledgements

We thank our Novartis colleagues: John Reece-Hoyes, Kushal Joshi, Qiong Wang and Dojna Shkoza for providing the cell lines; Walter Carbone, Judith Knehr for help with sequencing; Anthony Sonrel and Somesh Sai for discussion about the analytical approach; Jeremy Jenkins for scientific discussions.

## Footnotes

1 We use this nomenclature in order to be consistent with the definition in R. Note that there is an alternative nomenclature, which defines *α* = 1/*λ* as dispersion and is used in edgeR [73] and DESeq2 [74].

2 By size, we are referring to the actual distribution of the points in space, NOT the number of points in the cluster. For a Gaussian ellipsoid, size is parameterized by the covariance matrix.

3 Run time was estimated using the system.time() function in R. The time shown here refers to the full dataset (12000 cells). Analysis was run on 64-bit Intel(R) Xeon(R) CPU E7-4850 v2 @ 2.30GHz with 1TB of RAM in R 3.4.1 under Red Hat Enterprise Linux Server release 6.9 (Santiago). SC3 was run on 8 cores, all other methods on a single core.

## References

1. Macosko EZ, Basu A, Satija R, Nemesh J, Shekhar K, Goldman M, et al. Highly parallel genome-wide expression profiling of individual cells using nanoliter droplets. Cell. 2015;161:1202–14.

2. Klein AM, Mazutis L, Akartuna I, Tallapragada N, Veres A, Li V, et al. Droplet barcoding for single-cell transcriptomics applied to embryonic stem cells. Cell. 2015;161:1187–201.

3. Zheng GXY, Terry JM, Belgrader P, Ryvkin P, Bent ZW, Wilson R, et al. Massively parallel digital transcriptional profiling of single cells. Nat Commun [Internet]. Nature Publishing Group; 2017;8:14049. Available from: http://www.nature.com/doifinder/10.1038/ncomms14049

4. Svensson V, Vento-Tormo R, Teichmann SA. Exponential scaling of single-cell RNA-seq in the last decade. arXiv [Internet]. 2017; Available from: https://arxiv.org/ftp/arxiv/papers/1704/1704.01379.pdf%0Ahttp://arxiv.org/abs/1704.01379

5. Tang F, Barbacioru C, Wang Y, Nordman E, Lee C, Xu N, et al. mRNA-Seq whole-transcriptome analysis of a single cell. Nat Methods. 2009;6:377–82.

6. Rosenberg AB, Roco CM, Muscat RA, Kuchina A, Sample P, Yao Z, et al. Single-cell profiling of the developing mouse brain and spinal cord with split-pool barcoding. Science [Internet]. American Association for the Advancement of Science; 2018 [cited 2018 Mar 20];eaam8999. Available from: http://www.ncbi.nlm.nih.gov/pubmed/29545511

7. Cao J, Packer JS, Ramani V, Cusanovich DA, Huynh C, Daza R, et al. Comprehensive single-cell transcriptional profiling of a multicellular organism. Science [Internet]. American Association for the Advancement of Science; 2017 [cited 2018 Jan 24];357:661–7. Available from: http://www.ncbi.nlm.nih.gov/pubmed/28818938

8. Haber AL, Biton M, Rogel N, Herbst RH, Shekhar K, Smillie C, et al. A single-cell survey of the small intestinal epithelium. Nature. 2017;551:333–9.

9. Grün D, Lyubimova A, Kester L, Wiebrands K, Basak O, Sasaki N, et al. Single-cell messenger RNA sequencing reveals rare intestinal cell types. Nature. 2015;525:251–5.

10. Jiang L, Chen H, Pinello L, Yuan G-C. GiniClust: detecting rare cell types from single-cell gene expression data with Gini index. Genome Biol [Internet]. BioMed Central; 2016;17:144. Available from: http://genomebiology.biomedcentral.com/articles/10.1186/s13059-016-1010-4

11. Tirosh I, Izar B, Prakadan SM, Wadsworth MH, Treacy D, Trombetta JJ, et al. Dissecting the multicellular ecosystem of metastatic melanoma by single-cell RNA-seq. Science [Internet]. American Association for the Advancement of Science; 2016 [cited 2018 Apr 10];352:189–96. Available from: http://www.ncbi.nlm.nih.gov/pubmed/27124452

12. Villani A-C, Satija R, Reynolds G, Sarkizova S, Shekhar K, Fletcher J, et al. Single-cell RNA-seq reveals new types of human blood dendritic cells, monocytes, and progenitors. Science (80-) [Internet]. 2017;356:eaah4573. Available from: http://www.ncbi.nlm.nih.gov/pubmed/28428369

13. Shalek Alex K., Satija Rahul, Shuga Joe, Trombetta John J., Gennert Dave, Lu Diana, et al. Single-cell RNA-seq reveals dynamic paracrine control of cellular variation. Nature [Internet]. Nature Publishing Group, a division of Macmillan Publishers Limited. All Rights Reserved.; 2014;510:363–9. Available from: http://www.nature.com/nature/journal/v510/n7505/abs/nature13437.html#supplementary- information

14. Han X, Wang R, Zhou Y, Yuan G-C, Chen M, Correspondence GG, et al. Mapping the Mouse Cell Atlas by Microwell-Seq. Cell [Internet]. Elsevier Inc; 2018 [cited 2018 Apr 6];172:1091–107. Available from: https://doi.org/10.1016/j.cell.2018.02.001

15. Regev A, Teichmann SA, Lander ES, Amit I, Benoist C, Birney E, et al. The human cell atlas. Elife. 2017;6.

16. Kiselev VY, Kirschner K, Schaub MT, Andrews T, Yiu A, Chandra T, et al. SC3: Consensus clustering of single-cell RNA-seq data. Nat Methods. 2017;14:483–6.

17. Žurauskiene J, Yau C. pcaReduce: Hierarchical clustering of single cell transcriptional profiles. BMC Bioinformatics. 2016;17.

18. Setty M, Tadmor MD, Reich-Zeliger S, Angel O, Salame TM, Kathail P, et al. Wishbone identifies bifurcating developmental trajectories from single-cell data. Nat Biotechnol [Internet]. 30 2016;34:637–45. Available from: http://www.ncbi.nlm.nih.gov/pubmed/27136076

19. Qiu X, Mao Q, Tang Y, Wang L, Chawla R, Pliner HA, et al. Reversed graph embedding resolves complex single-cell trajectories. Nat Methods [Internet]. Nature Publishing Group; 2017;14:979–82. Available from: http://www.nature.com/doifinder/10.1038/nmeth.4402

20. Tsoucas D, Yuan G-C. GiniClust2: a cluster-aware, weighted ensemble clustering method for cell-type detection. Genome Biol [Internet]. BioMed Central; 2018 [cited 2018 May 14];19:58. Available from: https://genomebiology.biomedcentral.com/articles/10.1186/s13059-018-1431-3

21. Kharchenko Peter V, Silberstein Lev, Scadden David T. Bayesian approach to single-cell differential expression analysis. Nat Meth [Internet]. Nature Publishing Group, a division of Macmillan Publishers Limited. All Rights Reserved.; 2014;11:740–2. Available from: http://www.nature.com/nmeth/journal/v11/n7/abs/nmeth.2967.html#supplementary-information

22. Finak G, McDavid A, Yajima M, Deng J, Gersuk V, Shalek AK, et al. MAST: a flexible statistical framework for assessing transcriptional changes and characterizing heterogeneity in single-cell RNA sequencing data. Genome Biol [Internet]. BioMed Central; 2015;16:278. Available from: http://www.ncbi.nlm.nih.gov/pubmed/26653891

23. Korthauer KD, Chu LF, Newton MA, Li Y, Thomson J, Stewart R, et al. A statistical approach for identifying differential distributions in single-cell RNA-seq experiments. Genome Biol. 2016;17.

24. Johnson MB, Wang PP, Atabay KD, Murphy EA, Doan RN, Hecht JL, et al. Single-cell analysis reveals transcriptional heterogeneity of neural progenitors in human cortex. Nat Neurosci. 2015;18:637–46.

25. Camp JG, Badsha F, Florio M, Kanton S, Gerber T, Wilsch-Bräuninger M, et al. Human cerebral organoids recapitulate gene expression programs of fetal neocortex development. Proc Natl Acad Sci [Internet]. 2015;201520760. Available from: http://www.pnas.org/lookup/doi/10.1073/pnas.1520760112

26. Bardy C, Van Den Hurk M, Kakaradov B, Erwin JA, Jaeger BN, Hernandez R V., et al. Predicting the functional states of human iPSC-derived neurons with single-cell RNA-seq and electrophysiology. Mol Psychiatry. 2016;21:1573–88.

27. Handel AE, Chintawar S, Lalic T, Whiteley E, Vowles J, Giustacchini A, et al. Assessing similarity to primary tissue and cortical layer identity in induced pluripotent stem cell-derived cortical neurons through single-cell transcriptomics. Hum Mol Genet. 2016;25:989–1000.

28. R Development Core Team R. R: A Language and Environment for Statistical Computing. R Found. Stat. Comput. 2011.

29. Huber W, Carey VJ, Gentleman R, Anders S, Carlson M, Carvalho BS, et al. Orchestrating high-throughput genomic analysis with Bioconductor. Nat Methods. 2015;12:115–21.

30. McCarthy DJ, Campbell KR, Lun ATL, Wills QF. Scater: Pre-processing, quality control, normalization and visualization of single-cell RNA-seq data in R. Bioinformatics. 2017;33:1179–86.

31. ATL. Lun, Bach K, Marioni JC. Pooling across cells to normalize single-cell RNA sequencing data with many zero counts. Genome Biol [Internet]. BioMed Central; 2016;17:75. Available from: http://genomebiology.biomedcentral.com/articles/10.1186/s13059-016-0947-7

32. Vallejos CA, Risso D, Scialdone A, Dudoit S, Marioni JC. Normalizing single-cell RNA sequencing data: challenges and opportunities. Nat Methods [Internet]. Europe PMC Funders; 2017;14:565–71. Available from: http://www.ncbi.nlm.nih.gov/pubmed/28504683

33. Wilcoxon F. Individual Comparisons by Ranking Methods. Biometrics Bull. 1945;1:80.

34. Ritchie ME, Phipson B, Wu D, Hu Y, Law CW, Shi W, et al. limma powers differential expression analyses for RNA-sequencing and microarray studies. Nucleic Acids Res. 2015;43:e47.

35. Soneson C, Robinson MD. Bias, robustness and scalability in single-cell differential expression analysis. Nat Methods [Internet]. Nature Publishing Group; 2018 [cited 2018 Apr 17];15:255–61. Available from: http://www.nature.com/doifinder/10.1038/nmeth.4612

36. Brennecke P, Anders S, Kim JK, Kołodziejczyk AA, Zhang X, Proserpio V, et al. Accounting for technical noise in single-cell RNA-seq experiments. Nat Methods. Nature Publishing Group; 2013;10:1093–5.

37. Andrews TS, Hemberg M. Modelling dropouts for feature selection in scRNASeq experiments. bioRxiv [Internet]. Cold Spring Harbor Laboratory; 2017;65094. Available from: https://www.biorxiv.org/content/early/2017/05/25/065094

38. Langfelder P, Zhang B, Horvath S. Dynamic Tree Cut?: in-depth description, tests and applications. Bioinforamtics. 2007;1–12.

39. Fraley C, Raftery AE. Model-based Clustering, Discriminant Analysis and Density Estimation. J Am Stat Assoc. 2002;97:611–31.

40. Ester M, Kriegel HP, Sander J, Xu X. A Density-Based Algorithm for Discovering Clusters in Large Spatial Databases with Noise. Proc 2nd Int Conf Knowl Discov Data Min [Internet]. 1996;226–31. Available from: http://www.aaai.org/Papers/KDD/1996/KDD96-037.pdf”www.aaai.org/Papers/KDD/1996/KDD96-037.pdf

41. Stijn van Dongen. Graph Clustering by Flow Simulation. University of Utrecht; 2000.

42. Enright AJ, Van Dongen S, Ouzounis CA. An efficient algorithm for large-scale detection of protein families. Nucleic Acids Res [Internet]. 2002;30:1575–84. Available from: http://www.ncbi.nlm.nih.gov/pubmed/11917018

43. Mardia K, Kent J, Bibby J. Multivariate Analysis. London Acad Press. 1979;

44. Baglama J, Reichel L. Augmented Implicitly Restarted Lanczos Bidiagonalization Methods. SIAM J Sci Comput [Internet]. 2005;27:19–42. Available from: http://epubs.siam.org/doi/10.1137/04060593X

45. Hubert L, Arabie P. Comparing partitions. J Classif. 1985;2:193–218.

46. Grün D, Muraro MJ, Boisset J-C, Wiebrands K, Lyubimova A, Dharmadhikari G, et al. De Novo Prediction of Stem Cell Identity using Single-Cell Transcriptome Data. Cell Stem Cell. 2016;19:266–77.

47. Crow M, Paul A, Ballouz S, Huang ZJ, Gillis J. Characterizing the replicability of cell types defined by single cell RNA-sequencing data using MetaNeighbor. Nat Commun [Internet]. 2018 [cited 2018 Sep 10];9:884. Available from: http://www.nature.com/articles/s41467-018-03282-0

48. Kuhns MS, Badgandi HB. Piecing together the family portrait of TCR-CD3 complexes. Immunol Rev. 2012;250:120–43.

49. Nourani MR, Farajpour Z, Najafi A, Imani Fooladi AA. Trefoil Factor Family 1 Is Involved in Airway Remodeling of Mustard Lung. Iran J Allergy Asthma Immunol. 2016;15:275–82.

50. Prokopovic V, Popovic M, Andjelkovic U, Marsavelski A, Raskovic B, Gavrovic-Jankulovic M, et al. Isolation, biochemical characterization and anti-bacterial activity of BPIFA2 protein. Arch Oral Biol. Pergamon; 2014;59:302–9.

51. Kuijlaars J, Oyelami T, Diels A, Rohrbacher J, Versweyveld S, Meneghello G, et al. Sustained synchronized neuronal network activity in a human astrocyte co-culture system. Sci Rep [Internet]. Nature Publishing Group; 2016 [cited 2018 Dec 6];6:36529. Available from: http://www.nature.com/articles/srep36529

52. Pollen AA, Nowakowski TJ, Shuga J, Wang X, Leyrat AA, Lui JH, et al. Low-coverage single-cell mRNA sequencing reveals cellular heterogeneity and activated signaling pathways in developing cerebral cortex. Nat Biotechnol. 2014;32:1053–8.

53. Frotscher M. Cajal-Retzius cells, Reelin, and the formation of layers. Curr. Opin. Neurobiol. 1998. p. 570–5.

54. Nowakowski TJ, Bhaduri A, Pollen AA, Alvarado B, Mostajo-Radji MA, Di Lullo E, et al. Spatiotemporal gene expression trajectories reveal developmental hierarchies of the human cortex. Science (80-) [Internet]. 2017;358:1318–23. Available from: http://www.ncbi.nlm.nih.gov/pubmed/29217575

55. Meyer G, Perez-Garcia CG, Gleeson JG. Selective expression of doublecortin and LIS1 in developing human cortex suggests unique modes of neuronal movement. Cereb Cortex [Internet]. 2002;12:1225–36. Available from: http://www.ncbi.nlm.nih.gov/pubmed/12427674

56. Gonzalez-Gomez M, Meyer G. Dynamic expression of calretinin in embryonic and early fetal human cortex. Front Neuroanat. 2014;8:41.

57. Martinez-Galan JR, Moncho-Bogani J, Caminos E. Expression of Calcium-Binding Proteins in Layer 1 Reelin-Immunoreactive Cells during Rat and Mouse Neocortical Development. J Histochem Cytochem. 2014;62:60–9.

58. Molyneaux BJ, Arlotta P, Menezes JRL, Macklis JD. Neuronal subtype specification in the cerebral cortex. Nat Rev Neurosci [Internet]. Nature Publishing Group; 2007 [cited 2018 Apr 25];8:427–37. Available from: http://www.nature.com/articles/nrn2151

59. Rouillard AD, Gundersen GW, Fernandez NF, Wang Z, Monteiro CD, McDermott MG, et al. The harmonizome: a collection of processed datasets gathered to serve and mine knowledge about genes and proteins. Database (Oxford). 2016;2016.

60. Miller JA, Ding SL, Sunkin SM, Smith KA, Ng L, Szafer A, et al. Transcriptional landscape of the prenatal human brain. Nature. 2014;508:199–206.

61. Lun MP, Monuki ES, Lehtinen MK. Development and functions of the choroid plexus-cerebrospinal fluid system. Nat. Rev. Neurosci. 2015. p. 445–57.

62. Pollen AA, Nowakowski TJ, Chen J, Retallack H, Sandoval-Espinosa C, Nicholas CR, et al. Molecular Identity of Human Outer Radial Glia during Cortical Development. Cell [Internet]. 2015;163:55–67. Available from: http://linkinghub.elsevier.com/retrieve/pii/S0092867415011241

63. Cooper JA. Molecules and mechanisms that regulate multipolar migration in the intermediate zone. Front Cell Neurosci [Internet]. 2014;8. Available from: http://journal.frontiersin.org/article/10.3389/fncel.2014.00386/abstract

64. Chen G, Sima J, Jin M, Wang KY, Xue XJ, Zheng W, et al. Semaphorin-3A guides radial migration of cortical neurons during development. Nat Neurosci. 2008;11:36–44.

65. Priddle TH, Crow TJ. Protocadherin 11X/Y a Human-Specific Gene Pair: an Immunohistochemical Survey of Fetal and Adult Brains. Cereb Cortex [Internet]. Oxford University Press; 2013 [cited 2018 Apr 25];23:1933–41. Available from: https://academic.oup.com/cercor/article-lookup/doi/10.1093/cercor/bhs181

66. Pollen AA, Nowakowski TJ, Chen J, Retallack H, Sandoval-Espinosa C, Nicholas CR, et al. Molecular Identity of Human Outer Radial Glia during Cortical Development. Cell [Internet]. 2015;163:55–67. Available from: http://www.ncbi.nlm.nih.gov/pubmed/26406371

67. Lodato S, Molyneaux BJ, Zuccaro E, Goff LA, Chen HH, Yuan W, et al. Gene co-regulation by Fezf2 selects neurotransmitter identity and connectivity of corticospinal neurons. Nat Neurosci. 2014;17:1046–54.

68. Cunningham F, Amode MR, Barrell D, Beal K, Billis K, Brent S, et al. Ensembl 2015. Nucleic Acids Res. 2015;43:D662–9.

69. Schuierer S, Roma G. The exon quantification pipeline (EQP): A comprehensive approach to the quantification of gene, exon and junction expression from RNA-seq data. Nucleic Acids Res. 2016;44.

70. Bilican B, Livesey MR, Haghi G, Qiu J, Burr K, Siller R, et al. Physiological normoxia and absence of EGF is required for the long-term propagation of anterior neural precursors from human pluripotent cells. PLoS One. 2014;9.

71. Langfelder P, Zhang B, Horvath S. Defining clusters from a hierarchical cluster tree: The Dynamic Tree Cut package for R. Bioinformatics. 2008;24:719–20.

72. Campello RJGB, Moulavi D, Sander J. Density-Based Clustering Based on Hierarchical Density Estimates. Adv Knowl Discov Data Min [Internet]. 2013;160–72. Available from: http://link.springer.com/10.1007/978-3-642-37456-2_14

73. Robinson MD, McCarthy DJ, Smyth GK. edgeR: a Bioconductor package for differential expression analysis of digital gene expression data. Bioinformatics [Internet]. 2010;26:139–40. Available from: http://www.ncbi.nlm.nih.gov/pubmed/19910308%5CPMC2796818

74. Love MI, Anders S, Huber W. Differential analysis of count data - the DESeq2 package [Internet]. Genome Biol. 2014. Available from: http://biorxiv.org/lookup/doi/10.1101/002832%5Cnwhttp://dx.doi.org/10.1186/s13059-014-0550-8

