## Supplementary_Information for "CellSIUS provides sensitive and specific detection of rare cell populations from complex single cell RNA-seq data"

### Supplementary Figures and Tables

**Table S1 (separate file): Genes selected by different methods for feature selection.**

The file contains 3 sheets, one for each method tested. Each sheet contains the following columns:

- Gene_id: Ensembl gene identifier
- Symbol: Gene symbol
- Description: description of the gene function
- Pval (if applicable): p-value assigned by feature selection method
- Diff_to_fit: Difference from fitted trend. Each feature selection decides whether a feature is informative or not by testing for difference to a global trend. For HVG, this is a mean-variance trend, for NBDisp, a mean-dispersion trend and for NBDrop a mean-dropout trend.
- Rank: Assigned rank of the feature. For NBDrop and HVG, this is the order of diff_from_fit, from largest to smallest. For NBDisp, the order goes from smallest to largest.
- Mean_exprs: mean normalized expression [log2] of the feature across the whole dataset.
- Pct_variance: Percentage of variance attributed to ‘cell line’

3

**Table S2: Composition of full and subsampled cell line datasets**

| | Full dataset | | Subset 1 | | Subset 2 | |
|---|---|---|---|---|---|---|
| | ***Number*** | ***%*** | ***Number*** | ***%*** | ***Number*** | ***%*** |
| A549 | 1320 | 11.3 | 400 | 8.0 | 80 | 2.0 |
| H1437 | 1116 | 9.6 | 270 | 5.4 | 3 | 0.08 |
| HCT116 | 1743 | 14.9 | 1400 | 28.0 | 1599 | 40.1 |
| HEK293 | 2002 | 17.1 | 1600 | 32.0 | 2000 | 50.2 |
| IMR90 | 1039 | 8.9 | 500 | 10.0 | 100 | 2.5 |
| Jurkat | 962 | 8.2 | 100 | 2.0 | 6 | 0.15 |
| K562 | 1604 | 13.7 | 379 | 7.6 | 70 | 1.8 |
| Ramos | 1892 | 16.2 | 350 | 7.0 | 125 | 3.1 |
| ***TOTAL*** | **11678** | **100** | **4999** | **100** | **3983** | **100** |

**Table S3: Sequencing statistics and QC cutoffs per batch**

| | **Batch 1: IMR90, HCT116** | **Batch 2: HEK293, H1437** | **Batch 3: A549, Ramos** | **Batch 4: K562, Jurkat, Ramos** | **Neuronal differentiation** |
|---|---|---|---|---|---|
| Reads / cell [mean] | 37’756 | 33’589 | 38’602 | 36’236 | 59’660 |
| UMI / cell [min, 1$^{st}$ quartile, **median,** 3$^{rd}$ quartile, max] | 4569,14224,**17869,** 23196, 64864 | 4774, 9844, **12960**, 19469, 88332 | 4377, 9774, **13543**, 22310, 56086 | 1009, 8896, **16600**, 23824, 78355 | 2827, 5044, **6989**, 9474, 76642 |
| Genes / cell [min, 1$^{st}$ quartile, **median,** 3$^{rd}$ quartile, max] | 900, 3166, **3646,** 4200, 7344 | 1010, 2838, **3278**, 3870, 9211 | 1303, 2556, **3142**, 4004, 6608 | 168 ,2550, **3652**, 4357, 6852 | 1371, 2084, **2606**, 3135, 7131 |
| Filter thresholds | Min_UMI: 2^13<br>Min_genes: 2^11<br>Max_pct_mt: 10 | Min_UMI: 2^12<br>Min_genes: 2^10.5<br>Max_pct_mt: 10 | Min_UMI: 2^12<br>Min_genes: 2^10.5<br>Max_pct_mt: 10 | Min_UMI: 2^12<br>Min_genes: 2^10.5<br>Max_pct_mt: 10 | Min_UMI = 2^11.5<br>Min_genes = 2^10.5<br>Max_pct_mt = 10 |
| % cells removed by RNA content filter | 6 | < 1 | < 1 | 3 | 17 |
| % cells removed by mt genes filter | 4 | < 1 | < 1 | < 1 | 15 |
| Total % cells removed | 9 | < 1 | < 1 | 3 | < 1 |
| Number of genes x cells passing QC | 9452 x 2823 | 9858 x 3274 | 9290 x 3118 | 8727 x 2701 | 10351 x 4857 |
| Cell type assignment | IMR90: 1039<br>HCT116: 1743<br>Doublet: 38<br>Unassigned: 3 | H1437: 1116<br>HEK293: 2002<br>Doublet: 0<br>Unassigned: 156 | A549: 1320<br>Ramos: 1769<br>Doublet: 29<br>Unassigned: 0 | K562: 1606<br>Jurkat: 962<br>Ramos: 123<br>Doublet: 0<br>Unassigned: 0 | n.a. |

**Table S4 (separate file): DE analysis between subclusters and main clusters in the neuroscience dataset.**

The file contains 1 sheet per comparison. So far, there are two sheets.

- G.sub_1_vs_all_G: compares the G.sub_1 population to all other cells in the G cluster
- G.sub_2_vs_all_G: compares the G.sub_2 population to all other cells in the G cluster
- G.sub_3_vs_all_G: compares the G.sub_3 population to all other cells in the G cluster
- CR.sub_vs_all_CR: compares the CR.sub population to all other cells in the CR cluster
- NP.sub_vs_all_NP: compares the NP.sub population to all other cells in the NP cluster
- N.sub_1_vs_all_N: compares the N.sub_1 population to all other cells in the N cluster
- N.sub_2_vs_all_N: compares the N.sub_2 population to all other cells in the N cluster

Each sheet contains the following columns:

- Gene_id: Ensembl gene ID
- Mean_exprs: Mean expression [log2(normalized counts +1)] across the whole dataset
- Mean_in_subgroup: Mean expression in the respective subgroup
- Pval, adj_pval: p-value (Wilcoxon test), adj_pval is adjusted p-value (Benjamini-Hochberg)
- Log2fc: Fold change, calculated as the difference in mean[log2(normalized counts +1)]
- DE_flag: is TRUE if abs(log2fc) > 0.5 and adj_pval < 0.05
- Chr, symbol, eg, gene_biotype, description: Additional gene info (chromosome, gene symbol, entrez gene identifier, gene biotype, short description of gene function)

**Table S5: Media composition for the in-vitro differentiation of cortical excitatory neurons from human pluripotent stem cells in suspension**

| Media name | Composition |
|---|---|
| **Phase I** | Advanced DMEM/F12 (Gibco, 12634010), GlutaMax (Gibco, 35050061) 1% v/v, Pen/Strep (Gibco, 10378016) 1% v/v, N-acetyl-cysteine (Sigma, A9165) 500µM, Heparin (Sigma, 375795) 2 µg/mL, SB431542 (Tocris, 1614/1) 10 µM, LDN193189 (Tocris, 6053/10) 100nM, XAV939 (Tocris,3748/10) 2µM, N2 Supplement (Gibco, A1370701) 0.5% (v/v). |
| **Phase II** | Advanced DMEM/F12, GlutaMax 1% v/v, Pen/Strep 1% v/v, N-acetyl-cysteine 500µM, Heparin 2 µg/mL, N2 Supplement 0.5% v/v, B27 Supplement (Gibco, 17504044) 1% v/v, FGF2 10ng/mL (first 4 days)/2.5ng/mL (rest of PhII), LDN193189 100nM, CHIR99021 (Tocris, 4953/10) 20nM, Retinoic Acid (Sigma, R2625) 5nM. |
| **Phase III** | Advanced DMEM/F12, GlutaMax 1% v/v, Pen/Strep 1% v/v, Heparin 2 µg/mL, N2 Supplement 0.5% v/v, B27 Supplement 0.4 % v/v, Forskolin (Tocris, 1099/10) 10 µM, CaCl2 0.6 mM. |
| **Phase IV** | Advanced DMEM/F12, GlutaMax 1% v/v, Pen/Strep 1% v/v, Heparin 2 µg/mL, N2 Supplement 0.5% v/v, B27 Supplement 1.0 % v/v, Forskolin 10 µM, CaCl2 0.6mM, BDNF (Peprotech, AF-450-02) 5.0 ng/mL, GDNF (Peprotech, AF-450-10) 5.0 ng/mL. |

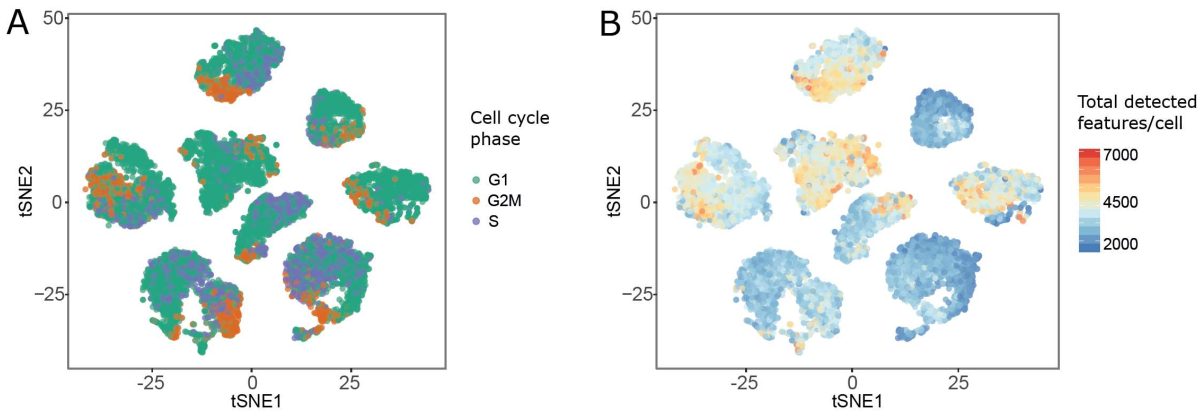


Figure S1: tSNE visualization of potential confounders in cell line dataset. A: tSNE-map, colored by predicted cell cycle phase. B: tSNE-map, colored by total detected features per cell.

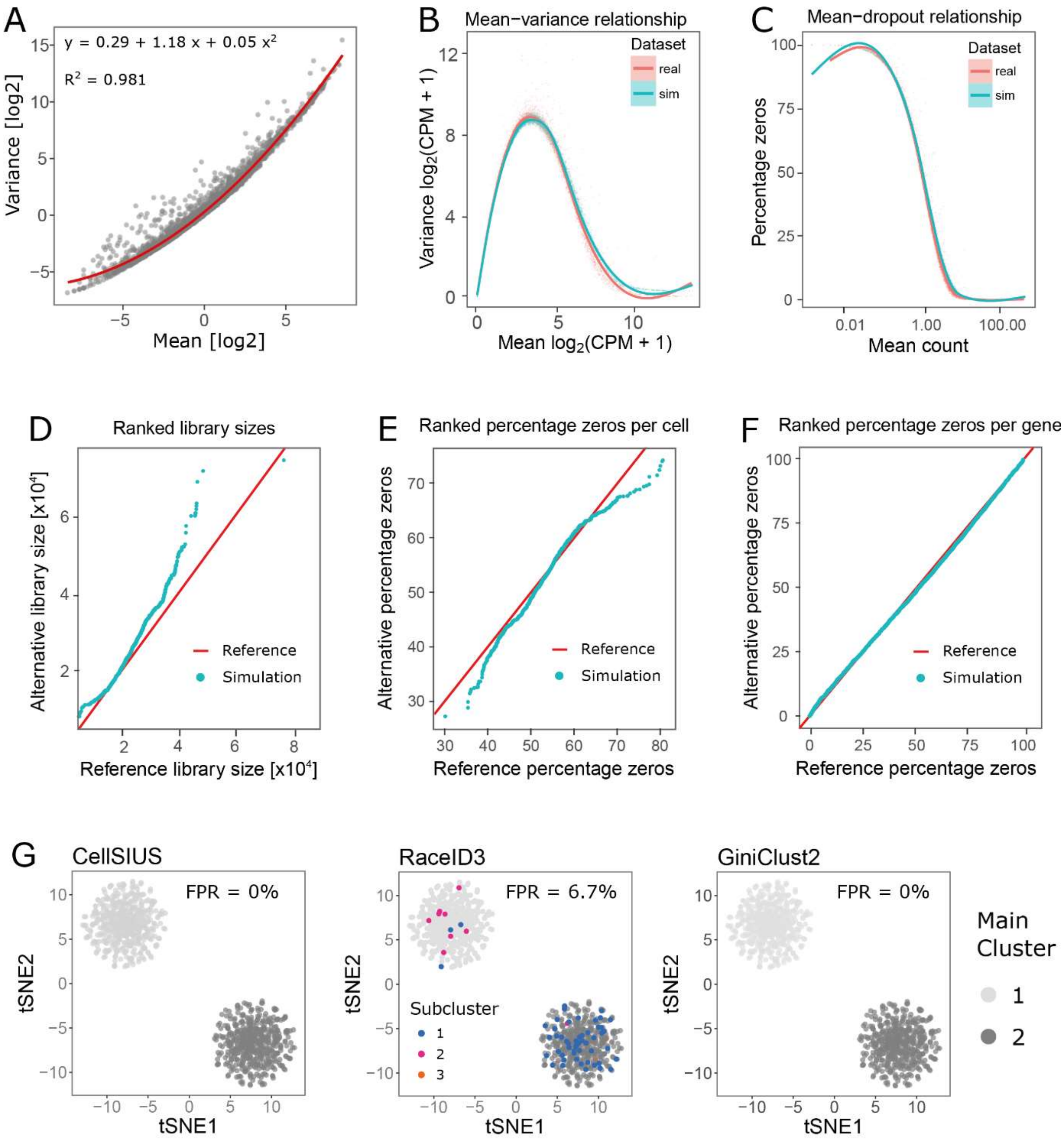


Figure S2: Generation of synthetic scRNA-seq data. A: Gene-wise mean-variance trend. The red line indicates a second order polynomial fit to the data that was used to estimate the expected variance as a function of the mean. B-F: Synthetic data (cyan) recapitulate the properties of experimental data (red). Plots show the relationship between the gene-wise mean and variance (B), the relationship between the mean count per gene and the fraction of zero counts / dropouts (C), ranked total number of counts per cell (D), ranked percentage of zero counts per cell (E) and ranked percentage of zeros per gene (F). G: Estimating false positive rates (FPR). The data shown are from two entirely homogeneous populations. CellSIUS and GiniClust do not falsely report rare cell types, whereas RaceID does.

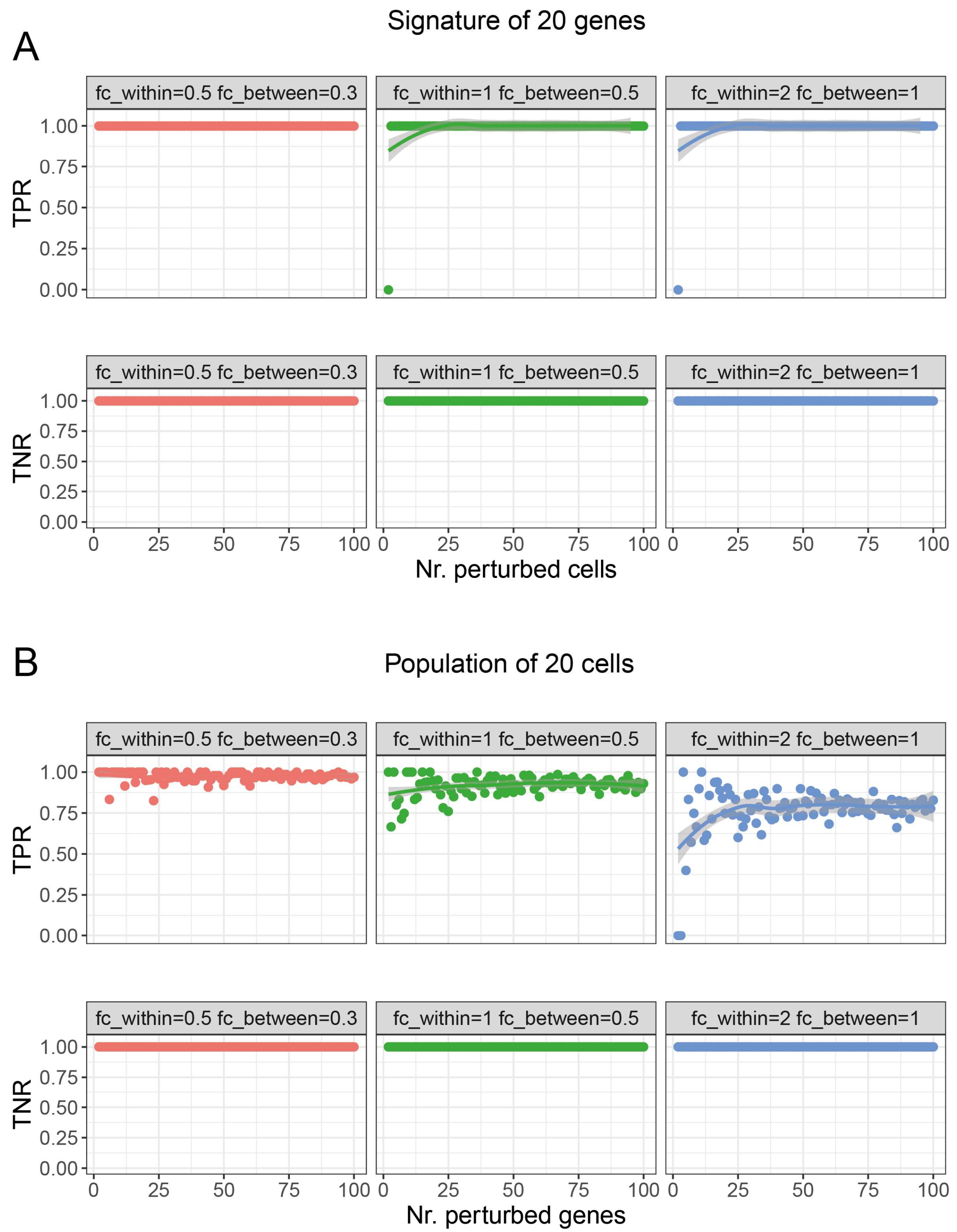


Figure S3: The recall or TPR (sensitivity) and TNR (specificity) of CellSIUS in detecting cell signatures of a number from 2 to 100 perturbed genes (A), and rare cells from 0.2% to 10% abundance (B), computed at different threshold parameters, fc_within (0.5, 1, 2) and fc_between (0.3, 0.5, 1).

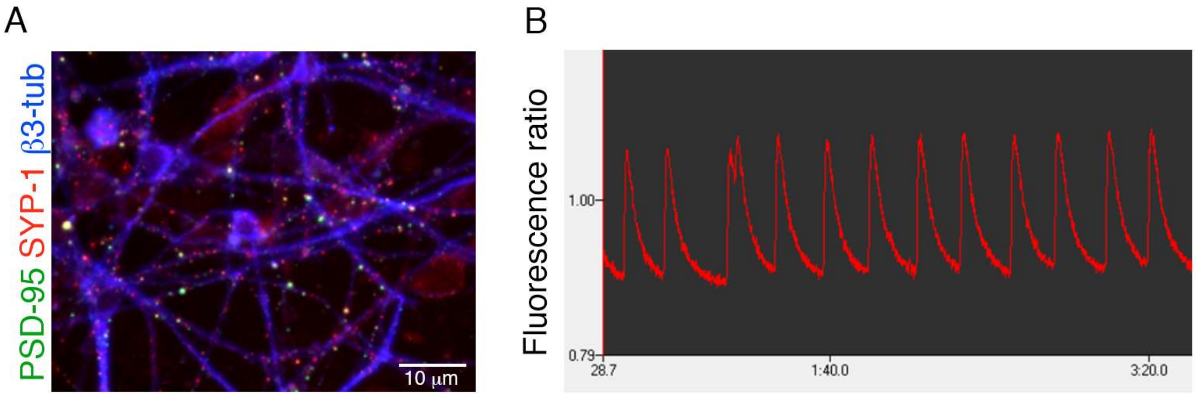


Figure S4: Maturation of hPSC-derived cortical neurons with rat glia co-culture. A: Immunohistochemical staining against post-synaptic density protein (PSD-95), Synaptophysin I (SYP-1) and β3-tubulin. B: Representative spontaneous calcium oscillation signal trace from a well of a high-density human cortical neuron / rat glia co-culture in 96-well format. Cells were loaded with the fluorescence calcium indicator FLIPR Calcium 6 dye (Molecular Probes) and observed for their spontaneous behaviour using the fluorescence plate reader FDSS7000EX (Hamamatsu).

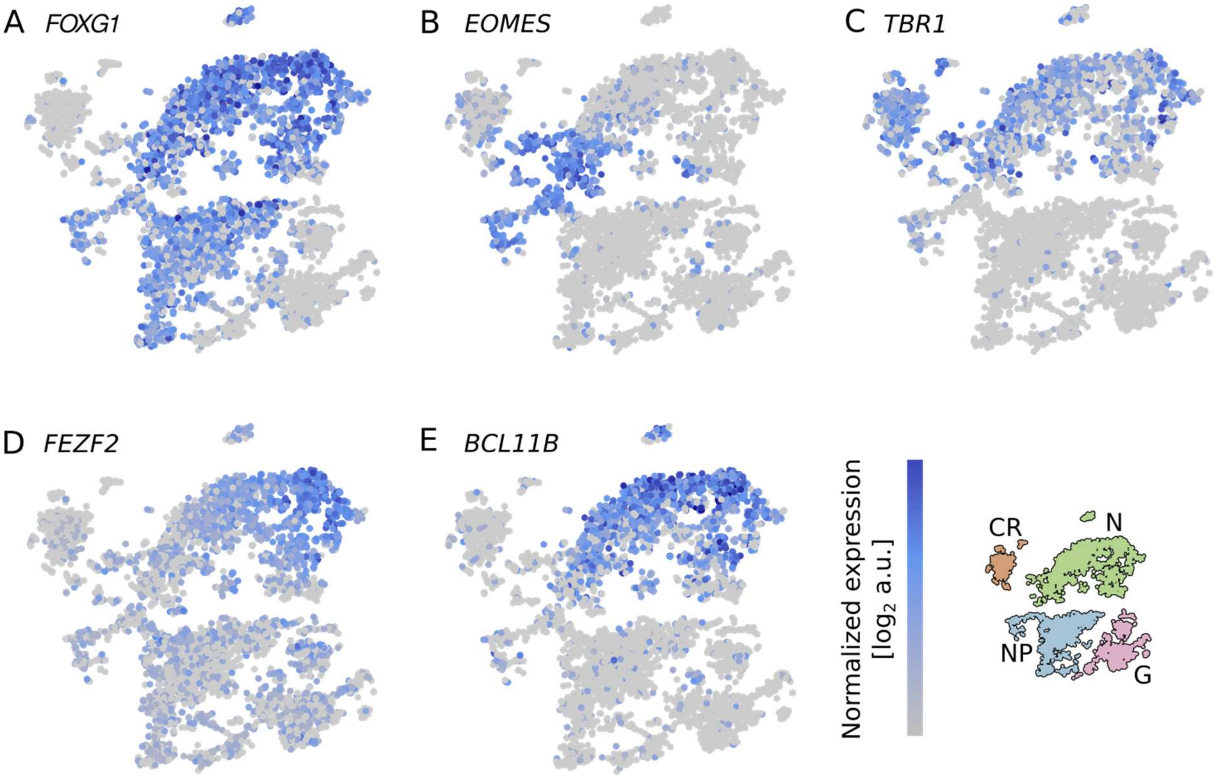


Figure S5: hPSC derived cortical neurons express characteristic marker genes. Shown are tSNE projections, colored by expression of A: FOXG1, B: EOMES, C: TBR1, D: FEZF2, E: BCL11B. The small map on the right shows the location of the main population on the tSNE projection (N: Neuron, NP: neuroepithelial progenitor, G: mixed glial cells, CR: Cajal-Retzius cells).

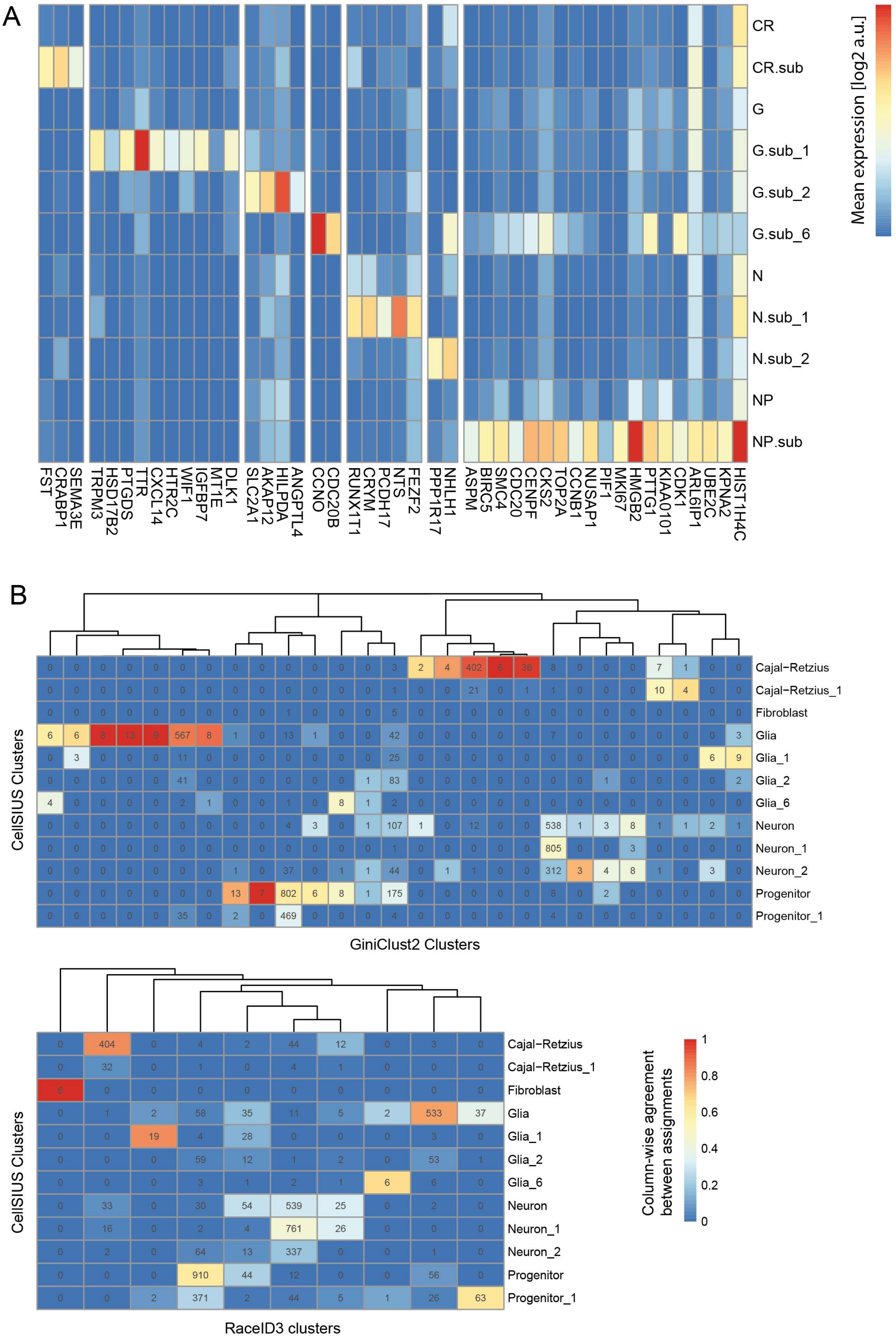
A
CR
CR.sub
G
G.sub_1
G.sub_2
G.sub_6
N
N.sub_1
N.sub_2
NP
NP.sub
Mean expression [log2 a.u.]
FST
CRABP1
SEMA3E
TRPM3
HSD17B2
PTGDS
TTR
CXCL14
HTR2C
WIF1
IGFBP7
MT1E
DLK1
SLC2A1
AKAP12
HILPDA
ANGPTL4
CCNO
CDC20B
RUNX1T1
CRYM
PCDH17
NTS
FEZF2
PPP1R17
NHLH1
ASPM
BIRC5
SMC4
CDC20
CENPF
CKS2
TOP2A
CCNB1
NUSAP1
PIF1
MKI67
HMGB2
PTTG1
KIAA0101
CDK1
ARL6IP1
UBE2C
KPNA2
HIST1H4C
B
Cajal−Retzius
Cajal−Retzius_1
Fibroblast
Glia
Glia_1
Glia_2
Glia_6
Neuron
Neuron_1
Neuron_2
Progenitor
Progenitor_1
CellSIUS Clusters
GiniClust2 Clusters
RaceID3 clusters
Column-wise agreement between assignments
1
0.8
0.6
0.4
0.2
0

Figure S6: Identification of cell subgroups in neuronal populations. A: Subgroups and their markers identified by CellSIUS. Rows correspond to clusters, columns to genes. Colors indicate mean expression level per cluster. B: Confusion matrix between cluster assignments by CellSIUS and Giniclust2 (top) and CellSIUS and RaceID3 (bottom). Rows correspond to CellSIUS assignment, columns to assignment by Giniclust2 and RaceID3, respectively. Numbers indicate the number of cells, colors indicate the degree of agreement, calculated per column.

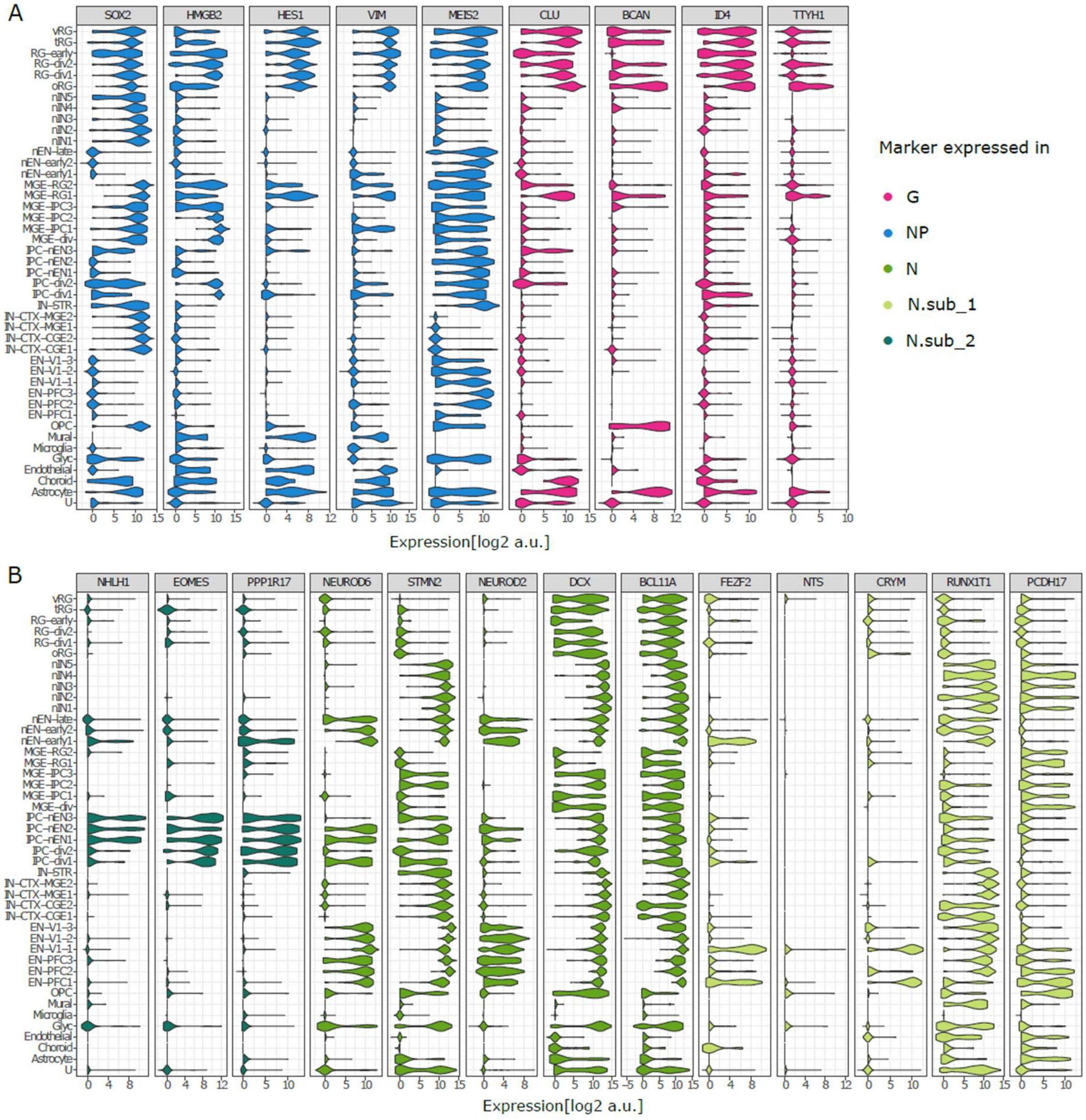


Figure S7: Comparison of neuronal population markers to scRNA-seq data from the developing human cortex. The data plotted are from a recent publication by Nowakowski *et al.* [1]. Shown are normalized expression values on a log2-scale, separated by cell type annotation as provided by the authors. For each gene, colors indicate which population expresses it highest in the *in vitro* corticogenesis model presented in this study. Abbreviations: RG = radial glia, vRG = ventricular RG, tRG = RG = radial glia, oRG = outer RG, IN = interneuron, nIN = newborn IN, EN = excitatory neuron, nEN = newborn excitatory neuron, MGE = medial ganglionic eminence, CGE = caudal ganglionic eminence, STR = striatum, CTX = cortex, V1 = primary visual cortex, PFC = prefrontal cortex, OPC = oligodendrocyte precursor cell, U = unknown.

#### List of supplementary data and code

The source of the CellSIUS R package: CellSIUS_1.0.0.tar.gz

R.code.tar.gz: compressed folder containing all the codes and processed data to reproduce the analysis discussed in the text. It includes:

- Supplementary_File_1_cell_line_data_setup.Rmd: Code for initial pre-processing of the cell line dataset.
- Supplementary_File_2A_cell_line_full.Rmd and Supplementary_File_2B_cell_line_full.html: Code (R Markdown) to reproduce the benchmarking on the full cell line dataset, and the generated html report.
- Supplementary_File_3A_cell_line_subset_1.Rmd and Supplementary_File_3B_cell_line_subset_1.html: Code (R Markdown) to reproduce the benchmarking on subset 1 of the cell line dataset, and the generated html report.
- Supplementary_File_4A_cell_line_subset_2.Rmd and Supplementary_File_4B_cell_line_subset_2.html: Code (R Markdown) to reproduce the benchmarking on subset 2 of the cell line dataset, and the generated html report.
- Supplementary_File_5A_rare_cell_identification_simdata.Rmd and Supplementary_File_5B_rare_cell_identification_simdata.html: Code (R markdown) to perform the benchmarking of the rare cell type identification methods on synthetic data, and the generated html report.
- Supplementary_File_6A_neuron_dataset_analysis.Rmd and Supplementary_File_6B_neuron_dataset_analysis.html: Code (R Markdown) to reproduce the analysis of the neuronal differentiation dataset.

- Supplementary_File_7_ K562_cleaned_counts.RData: processed count data used as a basis to generate simulated data as descried in supplementary file 5A,B.
- Supplementary_File_8_ sce_raw.RData: Count data and metadata for the cell line dataset, provided as an SCESet object used by the scater R package.
- Supplementary_File_9_ subset_1_idx.RData: Vector of column indices to subset Supplementary_File_8_ sce_raw.RData in order to generate subset 1 of the cell line dataset.
- Supplementary_File_10_ subset_2_idx.RData: Vector of column indices to subset Supplementary_File_8_ sce_raw.RData in order to generate subset 2 of the cell line dataset.
- Supplementary_File_9_benchmarking_incidence_subtlety.Rmd: Code to run the benchmarking on the in silico generated populations with different incidence and subtlety of transcriptional difference.

1. Nowakowski TJ, Bhaduri A, Pollen AA, Alvarado B, Mostajo-Radji MA, Di Lullo E, et al. Spatiotemporal gene expression trajectories reveal developmental hierarchies of the human cortex. Science (80- ) [Internet]. 2017;358:1318–23. Available from: http://www.ncbi.nlm.nih.gov/pubmed/29217575
